# Decision heuristics in contexts exploiting intrinsic skill

**DOI:** 10.1101/2022.04.01.486746

**Authors:** Neil M. Dundon, Jaron T. Colas, Neil Garrett, Viktoriya Babenko, Elizabeth Rizor, Dengxian Yang, Máirtín MacNamara, Linda Petzold, Scott T. Grafton

## Abstract

Heuristics can inform human decision making in complex environments through a reduction of computational requirements (accuracy-resource trade-off) and a robustness to overparameterisation (less-is-more). However, tasks capturing the efficiency of heuristics typically ignore action proficiency in determining rewards. The requisite movement parameterisation in sensorimotor control questions whether heuristics preserve efficiency when actions are nontrivial. We developed a novel action selection-execution task requiring joint optimisation of action selection and spatio-temporal skillful execution. Optimal choices could be determined by a simple spatial heuristic, or by more complex planning. Computational models of action selection parsimoniously distinguished human participants who adopted the heuristic from those using a more complex planning strategy. Broader comparative analyses then revealed that participants using the heuristic showed combined decisional (selection) and skill (execution) advantages, consistent with a less-is-more framework. In addition, the skill advantage of the heuristic group was predominantly in the core spatial features that also shaped their decision policy, evidence that the dimensions of information guiding action selection might be yoked to salient features in skill learning.

**Author Summary:** We often must choose between actions and then execute them, e.g., a tennis player chooses between a forehand and backhand and then skilfully executes the shot. To select actions, the player might plan their action with either shot, and select whichever simulated outcome is more advantageous. However, a more efficient approach might instead be to use a “heuristic”, i.e., a simpler rule, such as, forehand always on one side of the court, and backhand on the other. In this work, we look at whether styles of planning are related to physical skill performing actions, e.g., would a more skillful tennis player be more likely to use planning or a heuristic? We use a new task that requires people to choose and execute complex actions. Regarding choices, we use computational modeling to identify which people use some degree of planning, and which people use a simpler heuristic. Then, regarding action execution, we reveal that heuristic decision makers are in fact more skilled. However, they are not superiorly skilled in all aspects of performance, showing an advantage solely in the aspect of skill most closely linked to the information (spatial) they use for their heuristic. We therefore reveal the first ever evidence that a relation exists between the complexity of our action-related decisions and how broadly we learn associated motor behaviour.

## Introduction

In naturalistic settings, our cognitive architecture for making goal-oriented decisions typically resolves an ecological utility problem, integrating both extrinsic and intrinsic dynamics. Extrinsically, selected actions should maximise reward capture in line with a complex external state - a soccer player in possession of the ball must select the most rewarding action (shoot or pass) by incorporating such parameters as their location relative to the goalposts, availability of teammates, wind direction, readiness of the opposition goalkeeper, and so on. While the player might base their decision by planning and comparing outcomes across all possible actions, a high dimension external state likely favours some manner of decision heuristic, i.e., where actions are selected using a subset of all available external state information (e.g., if within 10 metres of the goalposts, shoot). Behavioural evidence verifies that a human decision policy can span different levels of planning complexity, with emerging neural evidence further suggesting that the brain harbours separate neural controllers for heuristics^1^.

The logic underscoring heuristic adoption is at least two-fold. Heuristics first offer a trade-off between accuracy and available resources. That is, where exhaustive planning might exceed computational resources or decision deadlines, heuristics offer a less laborious means to achieve a proxy for optimal action-selection policy^1, 2^. An alternative “less-is-more” rationale, inspired by machine learning principles, considers heuristics as the optimal means to avoid overfitting in uncertain environments. That is, in uncertain environments, a plan with too many parameters will likely pick up on stochastic noise and create more prediction errors across choices than a function that uses fewer parameters, even if the latter function produces a biased estimate^3^.

However, much like the areas of reinforcement learning and value-based decision making, evidence that humans exploit heuristics has emerged predominantly in contexts that do not consider intrinsic dynamics, such as skilled motor output, as a determining factor in reward yields. For example, in recent work^1^, simple button presses in a virtual task emulated foraging outcomes that probabilistically imparted a positive (partial increase), negative (partial decrease) or nonlinearly negative (complete erasure) impact on ongoing reward scores; human participants adopted a heuristic stimulus-driven policy that primarily avoided the nonlinear outcome, consistent with accuracy-resource trade-offs. Meanwhile, the less-is-more principle has been empirically supported in forecasting contexts such as weather^3^, investments^4^ and sporting events^5^. Simple- action probabilistic emulations and forecasting can innovatively replicate much of the extrinsic reward-oriented cognitive challenges presented by dynamic naturalistic environments, however, they probe only one side of the ecological utility dilemma. Lost in both paradigm formats are additional dynamic cost dimensions associated with effort^6, 7^, motor plasticity^8, 9^, and a broader sense of agency^10^, all of which integrate with external factors in the ultimate utility of selected actions in a momentary situation^11, 12^.

To our knowledge, no study has characterised heuristic adoption by humans when they select state- appropriate actions in selection-execution contexts, i.e., not only is there a correct action for a given state, but the proficiency of that selected action subsequently scales the level of reward and generates independent intrinsic error distributions such as spatial and temporal motor skill. According to sensorimotor control theory, such intrinsic error distributions are often attenuated by increasing the parameterisation of movement, e.g., by implementing forward-models or simulations^27–29^. This raises the question: are decision heuristics still efficient when actions are nontrivial; or does the requirement of a skilled physical action instead preserve the value of complex planning? We additionally do not know how decision heuristics relate to individual differences in skill. On the one hand, higher skill should improve both the time to generate, and the subsequent predictive utility of parametrising an impending action. In this case, higher-skilled individuals might be more likely to inform action choices with complex plans. However, an alternative prediction stems from the computational underpinnings of how motor learning evolves. Here, the commonly held view is that thorough deliberation dominates early in motor learning^13^, presumably while skill levels are also at their lowest. Thus, if the consequence of motor learning is a shift from low to high skill, in tandem with a shift from situational deliberation to less intensive (and more heuristic-like) draws from a cache of motor-memory strategies^13, 14^, higher-skilled individuals might be more inclined to inform action choices with heuristics.

To address these outstanding questions, we developed a novel task, where trialwise reward required joint optimisation of action selection (between two possible actions) and the subsequent execution of those actions (selection-execution task). We describe the task here in detail to provide the reader with an intuition for how it can identify participants using heuristics or more complex planning during action selection. Each trial involves selecting one of two computer cursors (that displace in different directions) to navigate from a starting position to a goal (start-goal, ‘SG’ pair) as efficiently as possible (see Figure1a-b). Reward on each trial depends on the level of fuel conserved, requiring the correct choice of cursor and thereafter skilful navigation in terms of both spatial skill and temporal control of non-linear acceleration (Figure 1d). SGs vary in terms of how well-suited they are to each cursor, with more ‘difficult’ choices therefore arising when an SG is similarly suited to both (Figure 2e).

**Figure 1.**
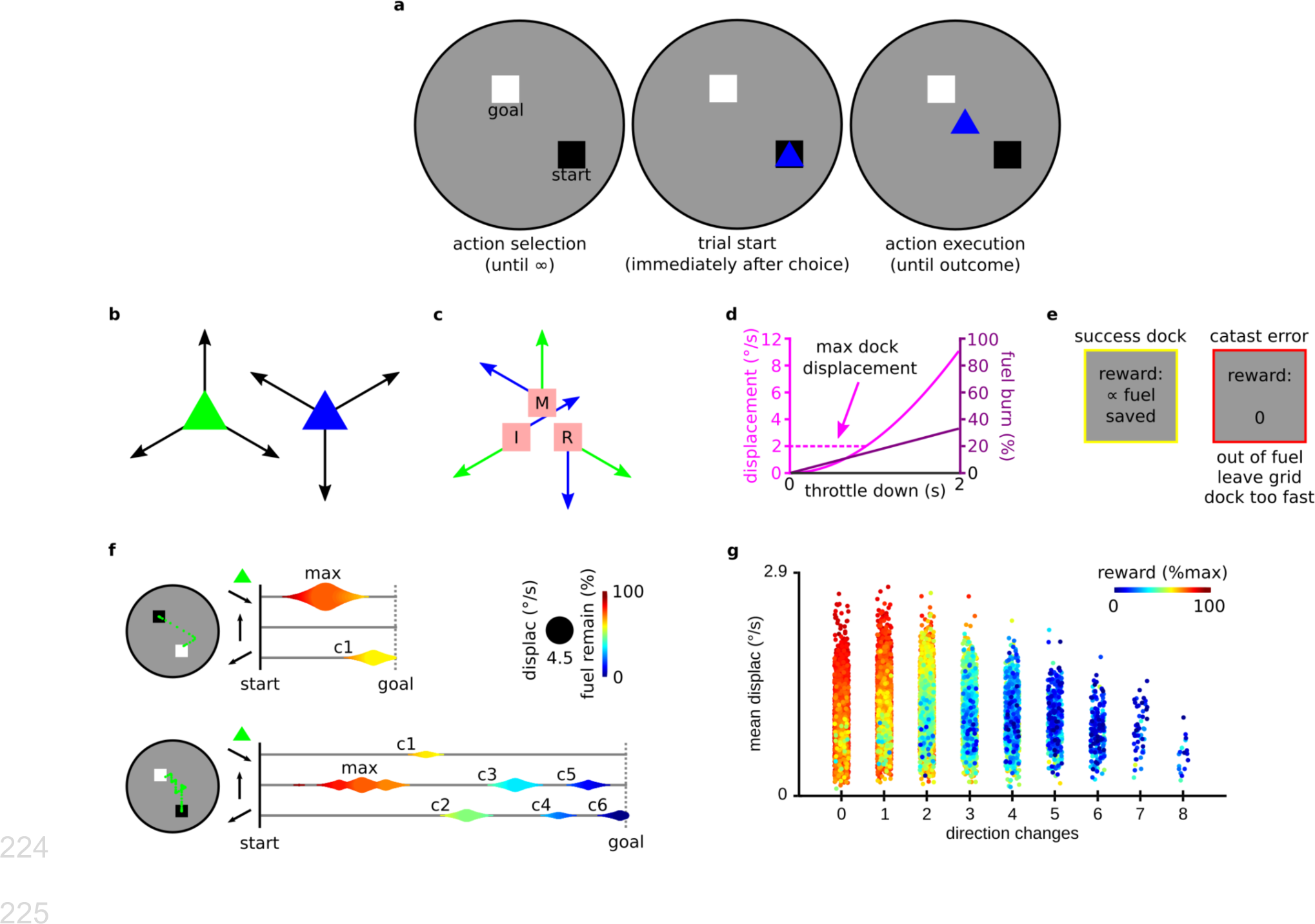
Action selection-execution task (boatdock) outline. **a,** on each trial, participants pilot one of two cursors between a start-goal pairing (SG). **b,** each cursor can accelerate in three unique directions. **c,** position of index (I), middle (M) and ring (R) finger of right hand on throttle buttons throughout the experiment, and cursor-specific throttle-vector mapping. Fuel burns any time a throttle is pushed down. Each trial allows six cumulative seconds of throttling before fuel depletes. **d**, throttle time linearly burns fuel, but nonlinearly increases displacement. Faster displacement is therefore more fuel efficient, however, a maximum dock displacement imparts additional temporal control requirements. **e,** successful docks yield a reward contingent on fuel conservation. This requires jointly maximising cursor choice for a given SG (action selection) in addition to spatial and temporal skill (action execution). Trials containing catastrophic errors - running out of fuel, leaving the grid, or docking above maximum displacement - yield no reward. **f,** schematic of two similar SGs with the same cursor but different performance dynamics. Three horizontal lines in each panel chart activity over time separately for each vector, while each vortex relates to a single throttle pulse. Top panel utilises fewer direction changes (marked with c1,…,cn), reaches a higher maximum displacement (depicted by diameter of largest vortex) and yields higher reward (depicted by colour). **g**, reward (depicted by colour), yielded on every successful trial across all participants (individual markers), is a joint function of spatial and temporal skill.

**Figure 2.**
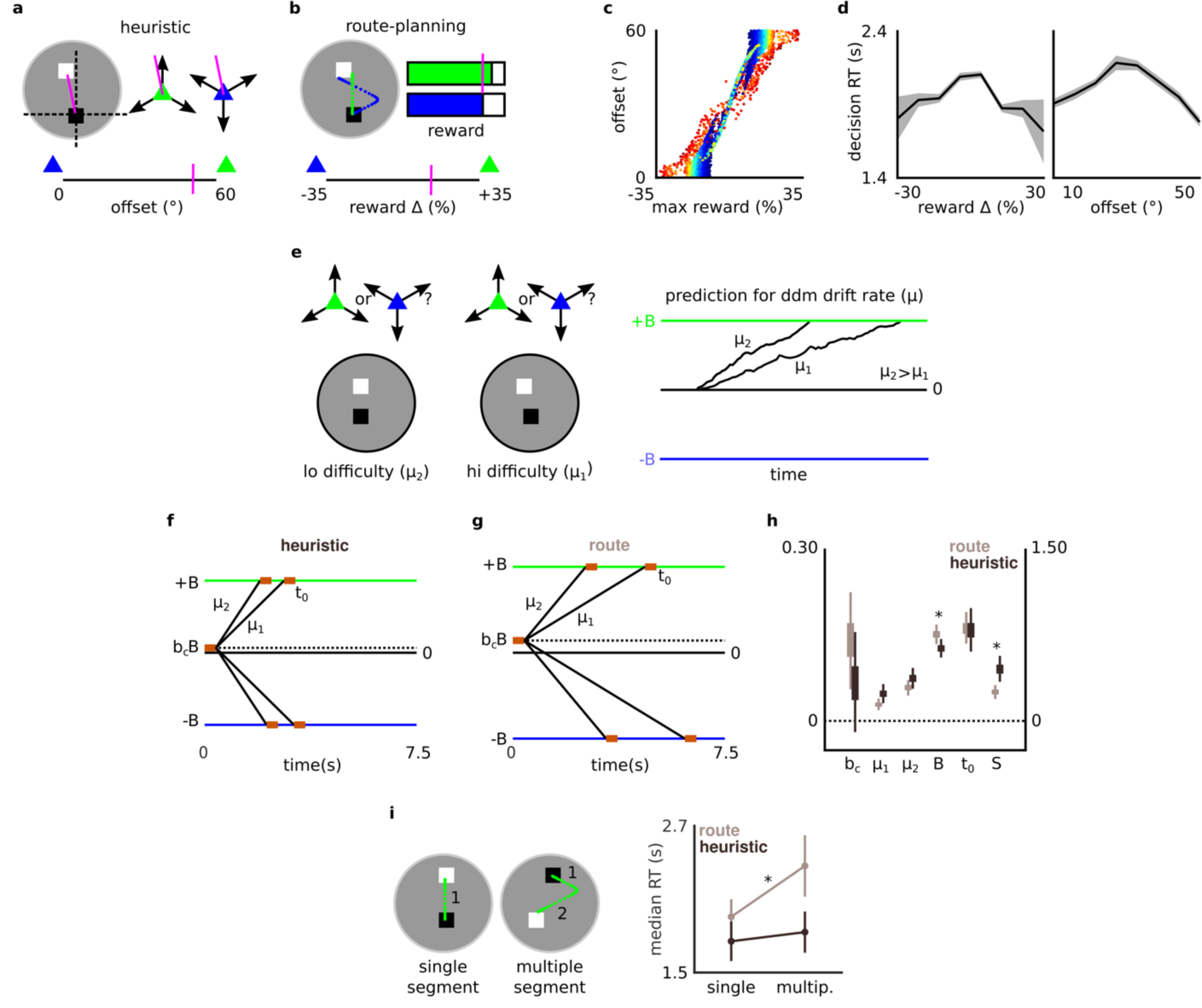
**Action-selection strategy identified by DDM framework. a-b**, a heuristic selects the cursor with a displacement vector with the least angular offset to the SG. Complex planning (route-planning) selects the cursor based on cursor- specific reward projections, i.e., incorporating additional spatial and/or temporal parameters into cursor evaluation over the heuristic. **c**, strategy-specific cursor suitability is imperfectly correlated across all trials from all participants. Hotter colours describe greater Euclidean distance between S and G. **d**, reaction time (RT) for action selection is greater on SGs where strategies ascribe equivalent suitability to both cursors. Solid black lines connect means of nine RT bins after sorting all participants’ trials by relevant strategy values. Gray shaded area depicts the standard error of the mean (S.E.M.) in each RT bin. **e**, DDM framework. A noisy evidence accumulation process terminates at a decision criterion (boundary). We hypothesised that difficulty arising in our task would modulate the rate of evidence accumulation (drift-rate *μ*). Depending on what strategy (heuristic or complex planning) a participant was using to make action selections, we’d see greater modulation of their drift-rate by difficulty arising from that strategy. For each participant’s choice and RT data, two target models allowed separate drift-rates *μ_1_* and *μ_2_* for high and low difficulty, respectively as per heuristic and route-planning strategy. Three groups of participants emerged, based on whether their data were best fitted by the heuristic (n=14), route-planning (n=19), or null (n=20) models. **f- g**, scaled schematic of DDM profile estimated for heuristic and route-planning (route) groups. **h**, comparison of DDM parameters between heuristic and route groups consistent with the latter integrating additional information into decision formation. Group’s differed in the sensitivity metric *S = (μ_1_+μ_2_)/(2B(1+|b_C_|))*, primarily due to route group having a credibly higher boundary (*B*). Route group’s bias (*b_c_*) was credibly above 0, indicating a bias away from the high-cost cursor. All parameters expressed in arbitrary units, except *t_0_*(in seconds). *t_0_* and *S* parameters are aligned with the right axis. Boxes and thin lines respectively represent the interquartile range (IQR) and highest density interval (HDI) of group- specific posteriors. (*0∉HDI(x_heuristic_-_route_)). **i**, group classifications verified independent of DDM parameters. Route group were uniquely slower to select the congruent cursor on routes requiring a multiple-segment route, where complex- planning demands likely become disproportionately greater than heuristic demands. (*p<0.01, Tukey-corrected). Vertical lines are S.E.M.

The suitability of a cursor to an SG can be guided by a simple heuristic that bases choice on a scalar value: the angle created by the SG and the nearest displacement vector of each cursor (“offset”, Figure 2a). Here, the more suitable cursor on a given SG is simply the one creating the smallest angular offset. Alternatively, participants might not use the angular heuristic alone and instead incorporate more complex planning into their action selection. Each cursor can travel in only three directions, meaning that cursors often need to perform some manner of segmented or “zig-zag” route to solve each SG. Thus, participants might have considered the impact of the required segmental paths when determining each cursor’s suitability. In other words, complex planning extends beyond the heuristic by additionally incorporating information such as the required action sequences, or predicted reward outcomes, into action selection (we operationalise complex planning more formally in the methods and results; see also Figure 2b).

We designed the task such that for each SG, the heuristic policy ultimately leads to the same recommended choice (selection) as any policy that also includes complex segmental planning, in order to ensure that both strategies result in similar action-execution requirements. However crucially, the degree of difficulty on each trial is different depending on whether a participant is using the heuristic or a complex planning policy. In other words, on some trials, an individual who is exploiting the heuristic will experience higher decision difficulty selecting a cursor than an individual using more complex planning and vice versa. Our computational approach to classifying participants as using either the heuristic or more complex planning centred on this strategy-specific emergence of choice difficulty, i.e., when a participant encountered decision difficulty. For example, if a participant showed increased signs of decision difficulty when the heuristic suggested more parity between cursors (i.e., similar angular offsets), to a greater extent than when a more complex plan suggested parity (i.e., routes with similar sequence requirements or reward outcomes), they were classified as likely using the heuristic.

To characterise decision difficulty emerging through qualitatively different planning styles, we incorporated the drift-diffusion model (DDM) into our computational modelling framework. The DDM, which has revealed comprehensive accounts of decision formation in both perceptual^15–18^ and goal-oriented^19–22^ contexts, formalises two-alternative action-selection deliberation as a gradual process of evidence accumulation toward one of two action-deterministic boundaries. The rate of evidence accumulation (also known as the “drift-rate”) is lower for more difficult decisions, such that easy decisions will likely have a much higher drift-rate than more difficult ones (see schematic in Figure 2e). In our analysis we exploited this relation between difficulty and drift-rate to identify participants’ strategies. I.e., if a participant is using the heuristic strategy, their data will be better fitted by a DDM that allows two drift-rates, one each for high and low difficulty trials, where difficulty is scored by the heuristic strategy. However, if a participant is using more complex planning, their data would instead be better fitted by a DDM that again allows two drift-rates, however for high and low difficulty trials as identified by more complex planning.

We therefore compared which of two DDMs (heuristic vs planning modulation of drift-rate) best fitted each participant’s data. To then verify our model classifications, we exploit the characteristics of a separate DDM parameter - boundary separation. This parameter enumerates the degree of evidence that an individual requires before executing a decision. Given that complex planning inherently requires more bits of information, we therefore expected that participants identified (by drift-rate modulations) as employing more complex planning would additionally show credibly larger boundary separation compared to those using the heuristic.

After classifying and verifying participants as likely using the heuristic or a more complex style of planning, we could then ultimately probe our core hypotheses regarding (i) whether heuristics are still efficient when selected actions are nontrivial and (ii) how decision heuristics relate to individual differences in skill. Regarding the initial question, we first tested whether participants using the heuristic required fewer runs of trials to persistently make state optimal action selections, i.e., select the more suitable cursor for a given SG, and whether they obtained higher overall level of reward in the task. We next tested the relation between heuristics and individual differences in skill and explored whether participants using the heuristic differed in terms of how skilfully they performed the action-execution portion of our selection-execution task. Specifically, we indexed action-execution skill using both spatial and temporal error dimensions, and tested whether both the overall level of these skills, in addition to their learning trajectories across the task, differed between participants using the heuristic and participants using complex planning.

## Results

Fifty-three healthy human participants performed 360 trials (six runs of 60) of a novel task framed as “boat docking” (Figure 1), in which reward yields require joint optimisation of action selection and action execution. On each trial, participants select one of two cursors to pilot between a randomly drawn start-goal pairing (SG; Figure 1a). Each cursor accelerates continuously in three unique directions (Figure 1b), burning fuel any time an accelerator button (throttle - Figure 1c) is down. One cursor imparts a higher motor execution cost via an incongruent key-mapping (Figure 1c). However, trialwise reward is contingent on fuel conservation, such that a selection policy that selects the cursor better suited to each SG, will yield higher reward. The two cursors accelerate with the same nonlinear function, and deplete fuel with the same linear function, i.e., faster displacement is more fuel efficient (Figure 1d). A maximum docking displacement rule (Figure 1d), imposing a speed limit on arrival, imparts additional temporal control demands. Thus, in addition to fewer direction changes (spatial error), greater temporal control maximises reward (Figures 1f-g). Finally, participants receive no reward for “catastrophic errors” (Figure 1e): when they run out of fuel, leave the grid, or attempt to dock above the maximum docking displacement.

### Summary behaviour - all participants

In general, participants followed task instructions and executed the task in a goal-oriented manner. In a series of separate summary Bayesian models we estimated the expected value (𝔼(x)) and highest density interval (HDI(x)) of group-level posteriors of key summary variables related to each participant’s task performance, that is, variables (such as median RT) that summarise across trials. At the group level, median reaction time to execute choices (median RT) was 1.70 s (𝔼(μ)=1.70, HDI(μ)=[1.46,1.935]). Participants registered catastrophic errors on 14.6 % of trials (𝔼(θ)=0.146, HDI(θ)=[0.141,0.151]). Participants displayed a preference (i.e., credibly above 50%) for the congruent cursor, selecting it on 62.3% of trials (𝔼(θ)=0.623, HDI(θ)=[0.616,0.630]), confirming that it imparted a lower cost relative to the incongruent cursor. However, notwithstanding this preference, participants were goal-oriented in their behaviour, selecting the cursor best suited to a given SG on 61.8% of trials (𝔼(θ)=0.618, HDI(θ)=[0.611,0.624]), i.e., above chance level (50%).

### Action selection using heuristic vs complex planning

To probe our core hypotheses we first needed to identify participants behaving in a goal-oriented manner and classify them on whether they were using the heuristic or more complex planning to inform action selection. In general, goal-oriented behaviour in our task requires participants to determine how suitable each cursor is for a given SG, during action selection (Figure 1B). To model how participants perform this evaluation, i.e., whether they likely used a heuristic or complex planning, we first scored each trial using two continuous measures. One measure enumerated the suitability of either cursor for its SG as derived by the heuristic, and the other as derived by complex planning. For the former, we designed our task such that the simplest policy to select suitable cursors for each SG involved the following heuristic: select the cursor with a vector subtending the smallest angular offset to that of the SG, i.e., incorporating only a single piece of spatial information into each choice (heuristic; Figure 2a). We therefore scored each trial using a continuous “offset” metric of how suitable either cursor was for its SG, as per this heuristic. This score ranged from 0 ° to 60 °, where 0 ° was a trial perfectly suited to the incongruent cursor and 60 ° was a trial perfectly suited to the congruent cursor. Thus, a participant using the heuristic to evaluate cursors would encounter maximal “difficulty” on SGs where this offset metric approached 30 ° (Figure 2a). Alternatively, participants had the option of incorporating additional complex action planning into their choices beyond the spatial heuristic. Such planning might require additional parameters over the heuristic, such as the spatial coordinates of simulated segmented routes, and/or the timing of pulses required in a simulated action sequence. In order to score each trial using a single continuous metric of how suitable either cursor was for its SG, as per more complex planning, we used a framework inspired by optimal control (“route-planningℍ; Figure 2b). At its optimum, complex planning in each SG would compare the route yielding the most reward with each cursor, and select the cursor with the highest value. In other words, projections of cursor-specific reward (i.e., conserved fuel) reflect a summary of planning that incorporates both spatial and temporal action parameters beyond the heuristic. We hypothesized that participants using more complex planning, thereby incorporating more bits of information into action selection beyond the heuristic (be they spatial or temporal), would accordingly show modulations during action selection that more-closely tracked cursor-specific reward projections than the heuristic score. We first used simulations (see Supplementary Materials: *Optimal route simulations*) to determine the routes yielding the most reward with each cursor for each SG, and the reward projected by each. Subtracting optimal incongruent reward from optimal congruent reward results in “route-planning” scores ranging from approximately -0.35 to 0.35 (where 1 corresponds to the starting fuel allocation on each trial). Thus, a participant selecting cursors using this strategy would encounter maximal “difficulty” on SGs with a route-planning score approaching 0. Note that the heuristic and route-planning scores always propose the same cursor for each SG, which ensures that action execution requirements are the same, regardless of which strategy a participant uses. However, importantly, the strategies are not perfectly correlated, due to additional nonlinear spatial and temporal information integrated into route-planning scores (Figure 2c). Across all participants, the reaction time (RT) for action selection was slower on more difficult trials, i.e., SGs where either strategy computed equivalent values for either cursor (heuristic=30 ° or route planning=0; Figure 2d), verifying that both methods for scoring SG successfully captured modifications to behaviour arising from increasing difficulty.

### DDM framework using drift-rate modulations to classify participants’ action-selection strategy

With each trial scored on one of two measures that could enumerate choice difficulty, we next fitted DDMs to (1) parse and later (2) verify the action-selection strategy of individual participants. The DDM (Figure 2e) describes a noisy sequential sampling process that accumulates evidence at an average “drift-rate” (*μ*) before reaching a decision criterion or “boundary” (*B* or *-B* for congruent or incongruent cursors, respectively). With a congruency bias *b_C_* favoring the congruent cursor where *b_C_* > 0, this process originates at a starting point *b_C_B*. Difficulty reduces the rate of evidence accumulation, which can be verified computationally when models containing two drift-rates (e.g., *μ_1_* and *μ_2_*), mapping respectively onto decisions presenting a high or low degree of difficulty, provide better model fits (Figure 2e). The first goal of our DDM framework was to distinguish people based on whether they used the heuristic or more complex planning to guide decisions (Figures 2a-b). We formally considered a participant to be using a specific strategy if their drift-rate was best modulated by difficulty arising from that strategy. To each participant’s set of trialwise choices and RTs we fitted three DDMs; a heuristic model, a route-planning model and a null model. Each model contained at least three free parameters: boundary *B*, congruency bias *b_C_*, and nondecision time *t_0_*. The null model was constrained such that *μ_1_* = *μ_2_* = 0, i.e., an arbitrary drift-rate indicating no deliberation over action selection. The heuristic and route-planning models had two free parameters, *μ_1_* and *μ_2_*, i.e., allowing two separate drift-rates that mapped respectively onto high or low difficulty trials. The heuristic model divided trials into equal bins reflecting high and low difficulty as calculated by offset scores. The route-planning divided trials into equal bins reflecting high and low difficulty as calculated by route-planning scores (see *Action selection using heuristic vs complex planning;* see Figures 2a-b). Based on model fits^24^ after adjusting for model complexity according to the Akaike information criterion with correction for finite sample size (AICc)^25, 26^, we confirmed that 14 participants’ choice and RT data were best fitted by the heuristic model, 19 participants’ data were best fitted by the route-planning model and 20 participants were best fitted by the null model, indicating neither strategy-specific difficulty modulated the rate of their evidence accumulation (Table 1). We hence refer to these three groups respectively as the “heuristic”, “route” and “nonplanner”.

**Table 1.**
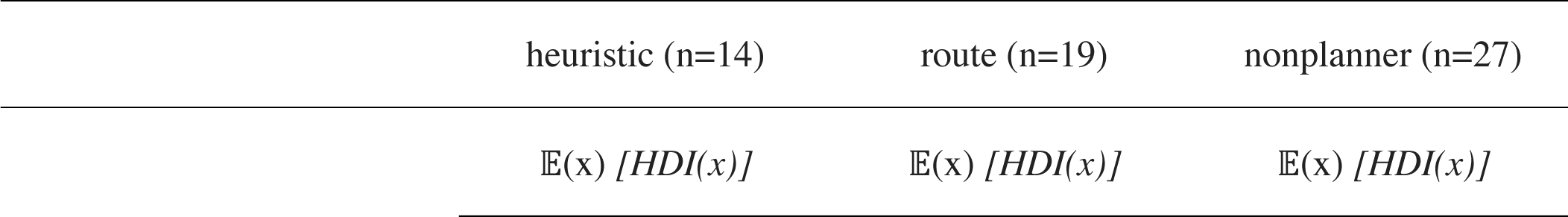
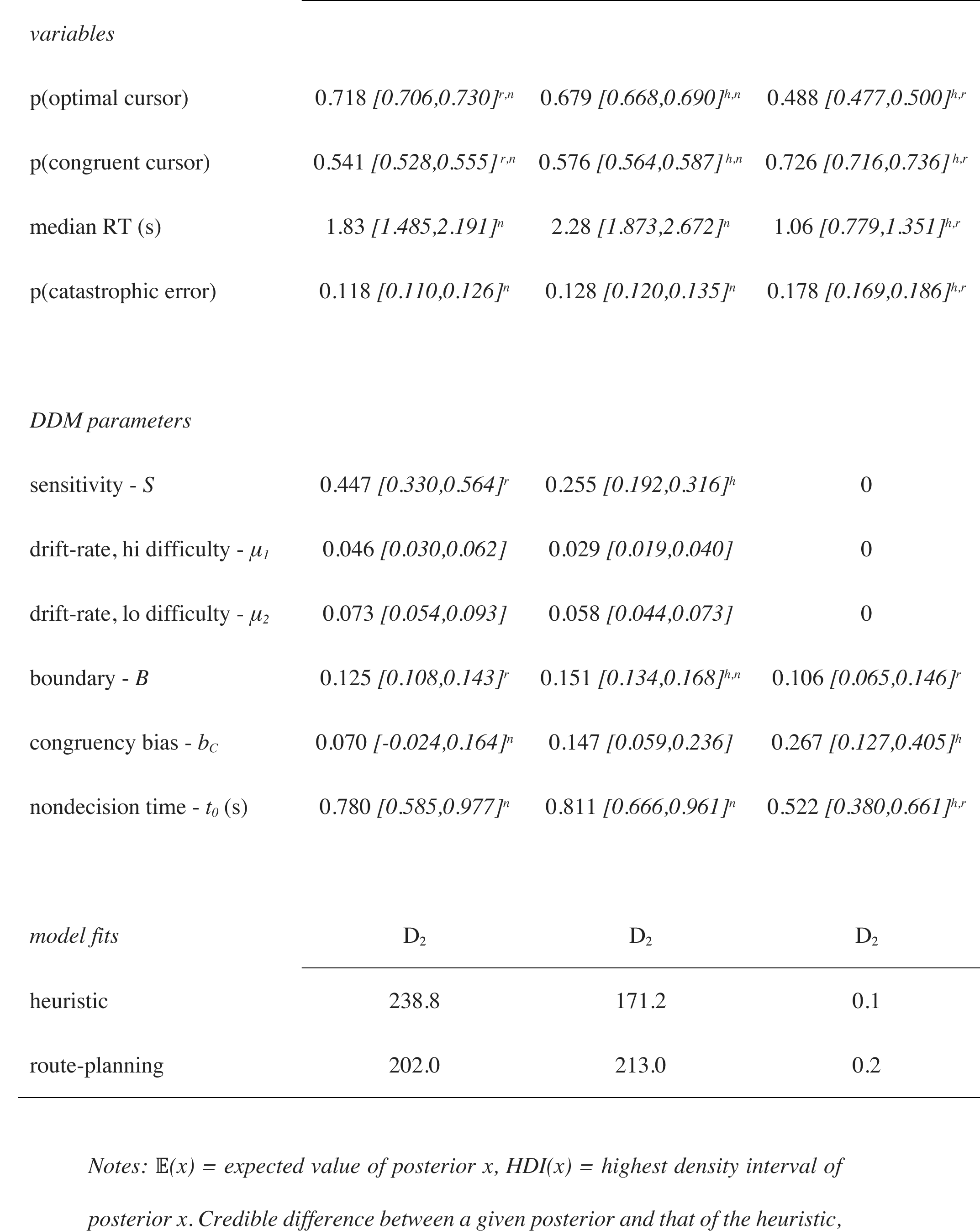
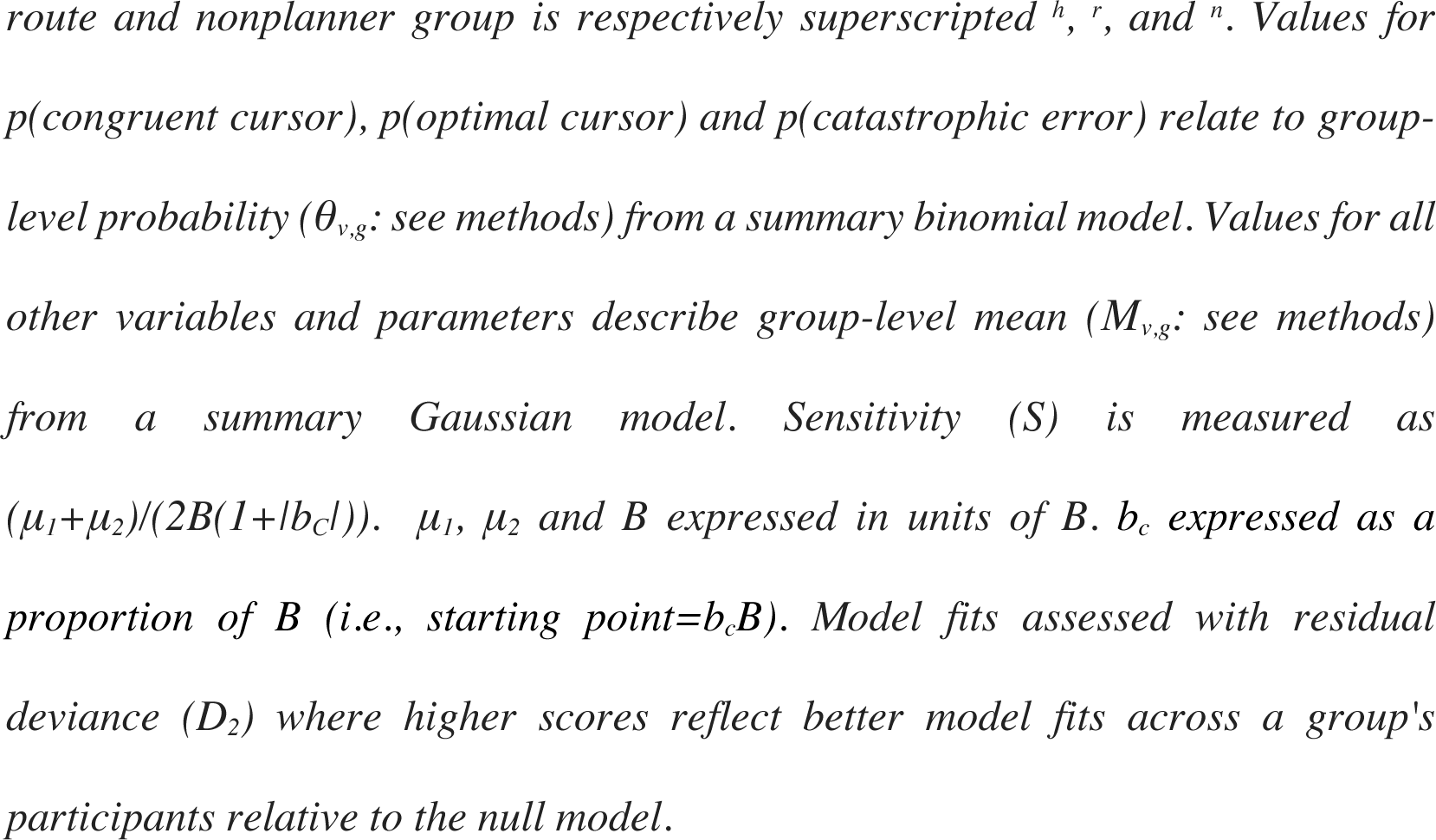
Group-specific posteriors for variables and DDM parameters and model fit scores

### Summary behaviour - group specific

We next looked at summary behaviour separately for each group, using Bayesian models that estimated the expected value (𝔼(x)) and highest density interval (HDI(x)) of posteriors that summarise group-specific estimates of key summary variables related to task. These group-specific estimates are presented in Table 1. Note that for the remainder of our results, anywhere we determined a difference between posteriors we required “credible evidence” (i.e., the highest density interval of one posterior minus the other to not subtend zero). Looking first at the differences between the heuristic and route groups, we observed that the heuristic group made the optimal choice on a credibly higher number (∼4%) of trials (𝔼(θ_heuristic_)=0.718, 𝔼(θ_route_)=0.679, 𝔼(θ_heuristic_-_route_)=0.039, HDI(θ_heuristic_-_route_)=[0.023,0.056]), which co-occurred with a credibly smaller preference for selecting the congruent cursor (𝔼(θ_heuristic_)=0.541, 𝔼(θ_route_)=0.576, 𝔼(θ_heuristic_-_route_)=- 0.035, HDI(θ_heuristic_-_route_)=[-0.017,-0.052]). However we observed no credible difference between these groups in terms of their median RT (𝔼(μ_heuristic_)=1.83, 𝔼(μ_route_)=2.28, 𝔼(μ_heuristic_-_route_)=-0.444, HDI(μ_heuristic_- _route_)=[-0.975,0.086]), nor their likelihood of registering a catastrophic error (𝔼(θ_heuristic_)=0.118, 𝔼(θ_route_)=0.128, 𝔼(θ_heuristic_-_route_)=-0.009, HDI(θ_heuristic_-_route_)=[-0.020, 0.001]). These initial summary comparisons suggest that in terms of discriminating the heuristic and route groups, choice optimality was a more telling behavioural feature than either the time needed to execute choices or frequency of catastrophic performance errors during the action-execution portion of each trial. Summary models further confirmed that the nonplanner group (i.e., the third group identified by our DDM as not performing any SG-appropriate deliberation) were not behaving in a goal-oriented manner as they did not credibly depart chance in terms of choice optimality (𝔼(θ)=0.488, HDI(θ)=[0.477,0.500])) and showed a strong bias toward the congruent cursor (𝔼(θ)=0.726, HDI(θ)=[0.716,0.736])). The remaining portion of the results will primarily focus on participants who behaved in a goal-oriented manner, i.e., we will compare and contrast the heuristic and route groups more thoroughly, in terms of DDM parameters, choice behaviour and skill. However, for completeness, we include data from the nonplanner group in Table 1, Figure 3 and Supplementary Table 1, and also include a supplementary section summarising their parametric and skill findings (see Supplementary Materials: *Nonplanner group*).

**Figure 3.**
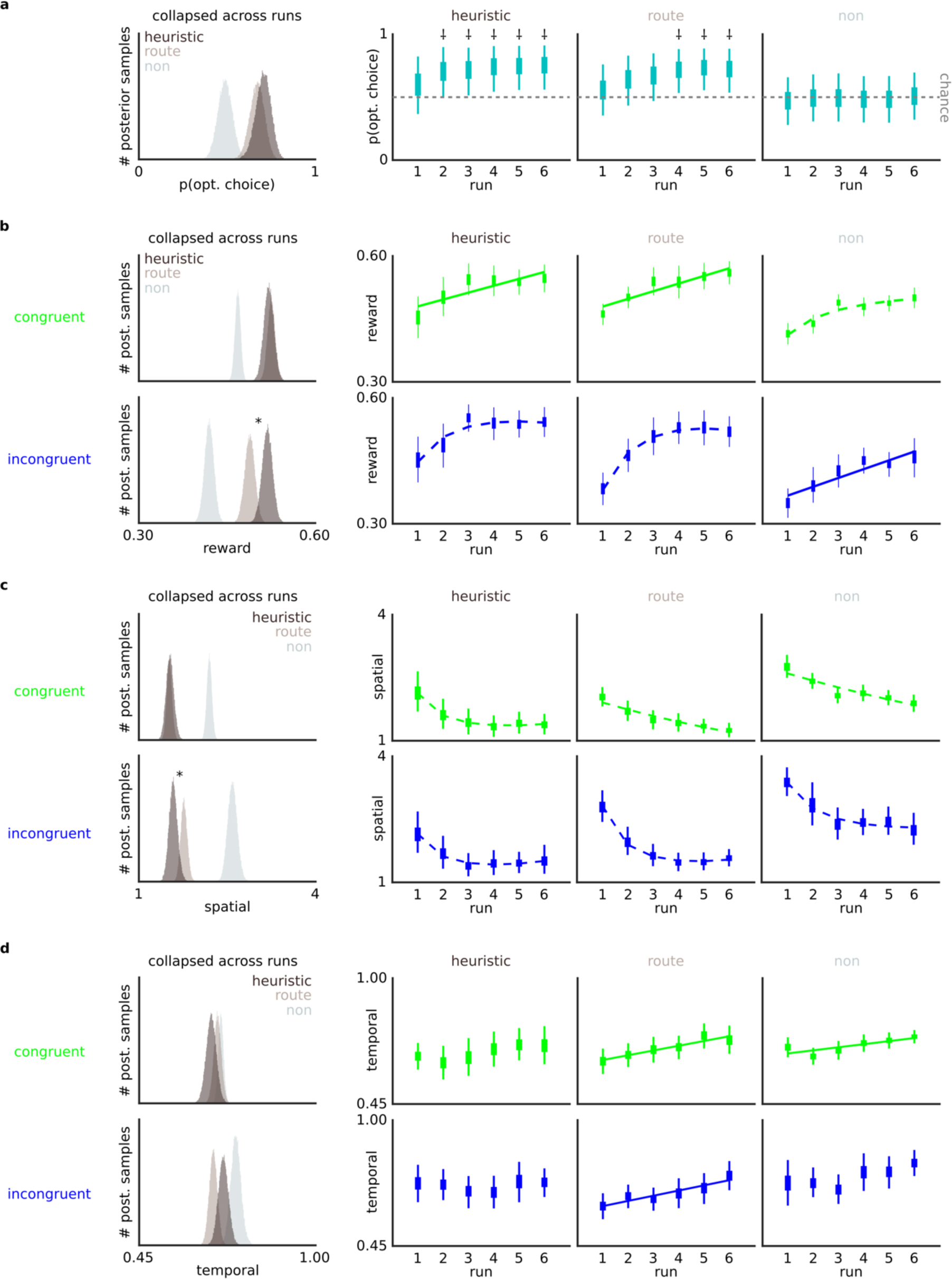
Heuristics group reach choice optimality more quickly and show a spatial-specific skill advantage. **a,** consistent with classic decision-heuristic models, low dimensional planning aligns with faster trajectories toward choice optimality. Hierarchical binomial model of choice behaviour demonstrates trade- off between the expediency and profundity of policy formation; heuristic group exceeded chance optimality by run 2, earlier than route group (run 4). ⸸ reflects runs where HDI of group-level θ posterior did not subtend 0.50, i.e., where group-level proportion of choices were credibly above chance optimality. **b-d** skill and skill learning suggest the dimensions of information guiding action selection are yoked to salient features in skill learning. Collapsing group-level posterior means across runs (skill) heuristic group yielded more reward with the high-cost cursor (**b**, histograms bottom panel), driven by superior spatial skill, i.e., the likely dominant feature in their action-selection policy (**c**, histograms bottom panel), with no route- heuristic difference in temporal skill (**d**, histograms). Asterisk relates to credible difference between route and heuristic groups, i.e., that the HDI of the deterministic distribution of their difference (heuristic-route) does not contain 0. Additionally, while route and heuristic group demonstrated skill learning in terms of reward and spatial skill (**b**-**c**, line plots), route group uniquely demonstrated learning in the temporal domain, a likely feature in their action-selection policy (**d**, line plots). Boxes and thin lines in line plots respectively represent IQR and HDI of hierarchical posteriors constraining individual-participant posteriors for a given measure, run and cursor. In both histograms and line plots, reward is the proportion of fuel preserved per trial (higher better), spatial is the number of direction changes (fewer better) and temporal is the distance-normalized difference between max and final velocity (higher better). Time-on-task (skill learning) effects estimated from deterministic regression models fitted across draws from each run’s posterior; credible (0 ∉ coefficient HDI) effects depicted by either a dashed (logarithmic) or solid (linear) line. Absence of any line reflects noncredible time-on-task effect.

### Verifying DDM classification of heuristic and route groups with DDM parameters

We next tested whether the group classifications ascribed by model fits based on strategy-specific drift-rate modulation were consistent with parameter values from the DDM. We specifically hypothesized that given complex planning’s incorporation of more bits of information into decisions, the route group would demonstrate larger boundary separation, i.e., consistent with them requiring more evidence to enact choices. In a single Bayesian model (see Methods) we estimated and compared group-specific means for each DDM parameter listed in Table 1. As an overall measure of performance, this model first compared the route and heuristic groups on the sensitivity metric *S=(μ_1_+μ_2_)/(2B(1+|b_C_|))*, which combines both drift-rates, congruency bias and the boundary. We observed credible evidence of group difference (𝔼(S_heuristic_)=0.447, 𝔼(θ_route_)=0.255, 𝔼(θ_heuristic_-_route_)=0.192, HDI(θ_heuristic_-_route_)=[0.056,0.322], Table1, Figure 2h). Lower sensitivity amongst the route group is consistent with their employed policy integrating more bits of information, as this metric is low when decision formation is jointly constrained by a low rate of evidence accumulation and a larger boundary (Figure 2g). Comparing each parameter separately, we observed that this effect was primarily driven by the degree of boundary separation, consistent with our hypothesis. The route group had a credibly higher boundary (*B*) than the heuristic group (𝔼(*B*_heuristic_)=0.125, 𝔼(*B*_route_)=0.151, 𝔼(*B*_heuristic_-_route_)=-0.025, HDI(θ_heuristic_-_route_)=[-0.050,-0.002], Table1, Figure 2h), but they did not have credibly different drift-rates, either for high difficulty SGs (𝔼(*μ_1_*_heuristic_)=0.046, 𝔼(*μ_1_*_route_)=0.029, 𝔼(*μ_1_*_heuristic_-_route_)=0.017, HDI(*μ_1_*_heuristic_-_route_)= [-0.002,0.036], Table1, Figure 2h) or for low difficulty SGs (𝔼(*μ_2_*_heuristic_)=0.073, 𝔼(*μ_2_*_route_)=0.058, 𝔼(*μ_2_*_heuristic_- _route_)=0.015, HDI(*μ_1_*_heuristic_-_route_)= [-0.001,0.040], Table1, Figure 2h). Also of note, we observed a credible bias toward the low cost action amongst the route group (𝔼(*b_c,_*_route_)=0.147, HDI(*b_c,_*_route_)=[0.059,0.236], Figure 2h), but not amongst the heuristic group (𝔼(*b_c,_*_heuristic_)=0.070, HDI(*b_c,_*_heuristic_)=[-0.024,0.164], Figure 2h). Together, these findings first provide parametric plausibility to the group classifications ascribed by our DDM, i.e., that the route group integrated more complex planning information into their choices. Secondly, the findings suggest that complex planning might be related to a stronger bias away from high-cost action, notwithstanding an overall tendency toward goal-oriented behaviour.

### Verifying DDM classification of heuristic and route groups with segmentation interaction

Our fundamental modelling assumption is that the route and heuristic groups are using qualitatively different planning strategies, and, for example, don’t simply differ in terms of general RT. In the above verification procedure we compared participants’ parameters that were fitted in the same DDM that also ascribed the group classifications (via strategy-specific drift-rate modulations). While the group difference in the boundary parameter was particularly consistent with our specific hypothesis predictions, we nonetheless sought additional evidence to verify the model classifications using data completely independent from the DDM. Specifically, we sought to identify moments in the task where the groups performed comparably in terms of RT (suggesting likely similarity in the computational requirements of their strategies), and other moments where their RT diverged (suggesting likely divergence in the computational requirements of their strategies). One such feature that we hypothesised would reveal such an interaction is the level of route segmentation in the optimal solution of SGs. In terms of information bits, the heuristic and complex planning should be most similar for SGs where the optimal solution is a single linear route from the start to the goal (single segment). However, for SGs where the optimal solution requires some kind of turn or zig-zag (multiple segment), complex-planning demands likely become disproportionately greater than heuristic demands, as the heuristic strategy can enumerate cursor-suitability using a single scalar value that is impervious to additional preparatory planning in action sequences. We therefore tested whether median RT showed an interaction between group (route, heuristic) and segmentation in SG solutions (single, multiple; see Figure 2i). Looking first at SGs that were better suited to the congruent cursor, a two-way mixed ANOVA of median RT returned a significant interaction between group and segmentation (F(1,31)=4.66;p=0.039; Figure 2i), underscored by a significant difference in median RT between single and multiple-segment routes for the route group (t(31)=4.04;p_tukey_=0.002) but not the heuristic group (t(31)=0.623;p_tukey_=0.924), consistent with our prediction. The same ANOVA on SGs better suited to the incongruent cursor returned no significant main effects or interactions (all p-values above 0.065). This absence of any effects in contexts suited to the high-cost action may reflect a greater level of noise interacting with the signal of action-selection processes, i.e., additional biases against using the incongruent cursor or additional preparatory processes related to incongruent finger-mapping. Importantly however, observing the predicted interaction at least in SGs suited to the congruent cursor reveals evidence independent of DDM parameters that the route and heuristic groups used qualitatively different planning strategies that differentially modulated RT depending on when their respective computational loads likely diverged.

### Are heuristics still efficient in selection-execution contexts?

After classifying and verifying participants as likely using the heuristic or a more complex style of planning, we next probed our core hypotheses, and first examined whether heuristics are efficient in a selection-execution context, i.e., when selecting nontrivial actions for execution. The remainder of the results section uses hierarchical Bayesian models that differ from the summary models above. The ensuing models allow for variance across trials with a hierarchy within the model itself. This hierarchical structure summarises trialwise measures within each participant with participant-level posteriors that were themselves constrained by a relevant hierarchical parameter. We further used separate hierarchical parameters depending on the hypothesis-relevant space of the model, e.g., group-by-run (see Methods for specific parameterisation). We first tested a prediction consistent with the ’less-is-more’ hypothesis, i.e., that people using heuristics would learn optimal policy more quickly. Specifically, we tested whether participants using the heuristic required fewer runs of trials to persistently make state optimal action selections. For this, we summarised choice optimality across time-on-task using a hierarchical binomial model. The model estimated the expected value (𝔼(x)) and highest density interval (HDI(x)) of choice optimality (θ) across a two-dimensional space described by group (heuristic, route, nonplanner) and run (1-6), i.e., group-specific choice optimality for each run (Figure 3a). Consistent with the less-is-more principle, the heuristic group demonstrated above-chance (0.50) choice optimality by the second run of trials (𝔼(θ_heuristic,run2_)=0.698, HDI(θ _heuristic,run2_)=[0.501, 0.898]), while in contrast, the route group’s choices did not credibly depart chance until the fourth run of trials (𝔼(θ_route,run4_)=0.712, HDI(θ _heuristic,run2_)=[0.533, 0.882]; Figure 3a, see Supplementary Table 1 for each group-by-run θ HDI). Thus, the heuristic group had a relative advantage of approximately 120 trials in reaching above-chance optimality with their choices. This finding corroborates the summary models above related to choice optimality, showing that heuristics are efficient in selection-execution contexts and possibly reflect a trade-off between how quickly a policy produces state-relevant choices, and the dimensionality of constituent planning.

Next, in terms of reward-specific value, we tested whether participants using the heuristic obtained a higher overall level of reward in the task. We first scored each trial in terms of reward, i.e., the proportion of the fuel tank conserved. We next summarised reward obtained across the task using a hierarchical Gaussian model. The model estimated the expected value (𝔼(x)) and highest density interval (HDI(x)) of average reward (μ) across a three-dimensional space described by group (heuristic, route, nonplanner), run (1-6), and cursor (congruent, incongruent), i.e., the credible ranges of group-specific reward, separately for each run, and separately again for each cursor. Figure 3b and Supplementary Table 1 contain each group-by-cursor-by-run μ estimate in addition to estimates collapsed across run. This model revealed that the heuristic group garnered higher reward yields across the task, however only with the high-cost (incongruent) cursor (Figure 3b). Merging posteriors across runs, the route and heuristic groups showed no credible differences in yielded reward (proportion of fuel conserved) using the congruent cursor (𝔼(μ_heuristic_)=0.550, 𝔼(μ_route_)=0.554, 𝔼(μ_heuristic_-_route_)=-0.004, HDI(μ_heuristic_-_route_)=[-0.025,0.017], Figure 3b), however the heuristic group amassed credibly higher yields (∼3% higher per trial) using the incongruent cursor (𝔼(μ_heuristic_)=0.548, 𝔼(μ_route_)=0.517, 𝔼(μ_heuristic_-_route_)=0.031, HDI(μ_heuristic_-_route_)=[0.006,0.055], Figure 3b). This reward advantage is an additional indication that heuristics are valuable in selection- execution contexts, consistent with the less-is-more principle. However, the cursor specificity of the effect additionally suggests that people employing the heuristic were not globally proficient across all aspects of the task, and instead had a reward advantage that was more prominent when employing high-cost action. This finding is additionally consistent with findings from our summary model above revealing the route planner group’s DDM bias away from selecting the congruent cursor.

### Comparisons of skill between route and heuristic groups

We have so far verified that a DDM framework parsimoniously distinguishes people on their likely use of heuristics or complex planning during action selection in a selection-execution task. Analyses regarding our first core hypothesis further suggest that those using the heuristic demonstrated advantages with respect to both decision policy formation and reward outcomes, consistent with the less-is-more principle. These analyses additionally suggest that action cost (i.e., dynamics involving the incongruent cursor) might be mediating the separation of these groups in some way. We next tested our second core hypothesis, i.e., how decision heuristics relate to individual differences in skill. We specifically tested two intrinsic skill dimensions underscoring reward outcomes during action execution - spatial and temporal skill. Spatial action execution was tracked by the number of direction changes on each trial, i.e., lower values reflect better performance on this measure which is modulated specifically by spatial precision. Temporal action execution was defined as the difference between the cursor’s maximum and final velocity (normalised by SG distance), i.e., higher values reflect better performance on this measure that indexes proficiency in temporal task demands requiring high max-velocities for more fuel-efficient displacement, while arriving at the goal below the maximum threshold (Figure 1d), and ideally as low as possible to further preserve fuel. We summarised performance in these two variables across the task using two separate hierarchical Bayesian models (Poisson for spatial skill, Gaussian for temporal skill), in each case using the same model space used above for reward. Each model estimated the expected value (𝔼(x)) and highest density interval (HDI(x)) of its relevant skill (μ) across a three-dimensional space described by group (heuristic, route, nonplanner), run (1-6), and cursor (congruent, incongruent), i.e., the credible ranges of group-specific skill, separately for each run, and separately again for each cursor. Figures 3c-d and Supplementary Table 1 contain each group-by-cursor-by-run μ estimate for each measure in addition to estimates collapsed across run. In terms of spatial skill, and consistent with our finding regarding reward advantage, the heuristic group again showed an advantage across the task, but only when using the incongruent cursor. Collapsing across runs, we observed no credible between-group difference with the congruent cursor (𝔼(μ_heuristic_)=1.53, 𝔼(μ_route_)=1.52, 𝔼(μ_heuristic_-_route_)=0.009, HDI(μ_heuristic_-_route_)=[-0.142,0.159], Figure 3c), but credibly fewer direction changes amongst the heuristic group with the incongruent cursor (𝔼(μ_heuristic_)=1.58, 𝔼(μ_route_)=1.76, 𝔼(μ_heuristic_-_route_)=-0.181, HDI(μ_heuristic_-_route_)=[-0.358,-0.011], Figure 3c). In terms of temporal skill, we observed no credible differences between the groups with either cursor. Collapsing across runs, we observed no between-group differences, with the congruent cursor (𝔼(μ_heuristic_)=0.671, 𝔼(μ_route_)=0.690, 𝔼(μ_heuristic_-_route_)=-0.020, HDI(μ_heuristic_-_route_)=[- 0.056,-0.018], Figure 3d) or incongruent cursor (𝔼(μ_heuristic_)=0.710, 𝔼(μ_route_)=0.677, 𝔼(μ_heuristic_- _route_)=0.031, HDI(μ_heuristic_-_route_)=[-0.007,0.068], Figure 3d). Given that the heuristic group reached state optimal choices more quickly (Figure 3a; Supplementary Table 1), i.e., they performed a higher volume of trials where their cursor selection theoretically reduced the need for direction changes, we re-ran the spatial skill model with trialwise direction changes adjusted by the optimal solution for the cursor selected for each given trial (i.e., observed direction changes - ideal direction changes). This model (see: Supplementary Materials - *Hierarchical Poisson with choice- normalised spatial skill*) returned identical results, confirming that notwithstanding their better choices, the heuristic group independently demonstrated greater spatial skill while piloting the incongruent cursor. This finding informs our second core hypothesis by linking heuristics to higher levels of skill. However, the spatial-specific nature of the heuristic group’s skill advantage further suggests that heuristics might not be related to higher skill proficiency in a global sense, but instead related to selective advantage in a reduced set of dimensions. Given its spatial nature, it raises the question as to whether their selective advantage in action execution might additionally be related to their action-selection strategy; i..e, the heuristic contains primarily spatial information, while more complex planning likely included additional dimensions (i.e., temporal) information.

### Time-on-task effects - skill learning

The previous section focused on group-by-cursor differences in skill when collapsing across runs, i.e., overall task performance. We next probed an additional component of our second core hypothesis and tested whether the route and heuristic groups also differed in terms of time-on-task trajectories in cursor-specific action execution (skill learning). For this, we took the uncollapsed (i..e, separate run-by-run) posteriors from each group-by-cursor dyad in the above hierarchical models of reward, spatial skill and temporal skill and drew samples to perform deterministic regression models. These regression models tested whether these features evolved over the task in either a linear (β_lin_) or logarithmic (β_log_) fashion, separately for each group and cursor (linear and nonlinear time-on-task effects). Figures 3b-d depict whether skill evolved across features in a linear or logarithmic fashion, while each group-by-cursor time-on-task coefficient (linear and logarithmic) is included in Supplementary Table 1. These skill learning models revealed additional divergence between the route and heuristic groups. The route group’s skill learning encompassed a broad range of features, i.e., they showed either linear or logarithmic improvement with each cursor for reward (cong: 𝔼(β_lin_)=0.160, HDI(β_lin_)=[0.103,0.218]; incong: 𝔼(β_log_)=0.564, HDI=[0.236,0.897]; Figure 3b), spatial skill (cong: 𝔼(β_lin_)=-0.157, HDI 𝔼(β_lin_)=[-0.211,-0.105]; incong: 𝔼(β_log_)=-0.499, HDI=[-0.728,-0.262]; Figure 3c) and temporal skill (cong: 𝔼(β_lin_)=0.124, HDI=[0.043 0.205]; incong: 𝔼(β_lin_)=0.134, HDI=[0.048,0.217]; Figure 3d). In contrast, the heuristic group only showed strong evidence of skill learning for reward (cong: 𝔼(β_lin_)=0.144, HDI(β_lin_)=[0.058,0.230]; incong: 𝔼(β_log_)=0.379, HDI(β_log_)=[0.008,0.763]; Figure 3b) and spatial skill (cong: 𝔼(β_log_)=-0.369, HDI(β_log_)=[-0.703,-0.024]; incong: 𝔼(β_log_)=-0.392, HDI(β_log_)=[-0.744,- 0.035]; Figure 3c), i.e., they demonstrated no credible skill learning with either cursor in the realm of temporal skill (Figure 3d; all β_lin_ and β_log_ HDIs subtend 0; see Supplementary Table 1). To corroborate this null result regarding temporal skill, we conducted a follow-up analysis. In summary, this nonparametric model compared how many individual participants in each group showed either linear or logarithmic time-on-task effects with temporal skill, i.e., a binomial design that could confirm different rates of skill learning between groups. We then performed a summary binomial model that computed group-specific proportions (θ) of participants that improved in temporal skill. This model confirmed that a higher number of participants in the route group improved relative to the heuristic group, both using the congruent cursor (10/19 vs 1/14;HDI(θ_route-heuristic_)=[0.145,0.647]) and using the incongruent cursor (10/19 vs 0/14;HDI(θ_route-heuristic_)=[0.068,0.600]). This additional analysis confirmed that the route and heuristic groups diverged in the feature of temporal skill learning, with less evolution in temporal skill demonstrated by the latter group. This finding further informs our second core hypothesis investigating the relation between decision heuristics and individual differences in skill, and further supports the idea that skill dimensions underscoring action execution might be related to a person’s action-selection strategy. Specifically, these skill-learning models suggest participants using the primarily spatial heuristic showed less learning across the task in the temporal domain. The route group, who in contrast were more likely to be incorporating temporal information into action selection, demonstrated credible learning in this dimension. Together with the above findings relating to the heuristic group’s overall superiority in spatial skill, these findings suggest that a yoked dimensionality might exist between planning and skill, i.e., in complex states, the number of features relevant (or not) for action-selection policy may predict the number of features most likely to undergo learning (or not) during action execution.

## Discussion

At least two schools of thought lend plausibility to the idea that humans might achieve optimal long-term ecological yields by basing goal-oriented decisions on subsets of information available in complex states. On the one hand, if either time or computational resources are restricted, humans might pragmatically trade off state-optimal parameterisation for reduced processing requirements (accuracy-resource trade-off). On the other hand, decisions informed by fewer parameters are more robust to the influences of misleading stochastic noise (less-is-more principle). In either case, extant knowledge on decision heuristics stems predominantly from action-trivial tasks that obviate intrinsic skill proficiency in determining behavioural outcomes. The present work directly addressed this shortcoming. We developed a novel selection-execution task requiring joint optimisation of action selection (of state-appropriate low-cost or high-cost cursors) and action execution (controlling cursors proficiently). Focusing first on action selection, cursor-state suitability could be determined by either a simple spatial heuristic strategy or more complex planning involving additional (spatial or temporal) action parameters. Using a between-group DDM framework, that exploited strategy-specific modulation of the drift-rate (evidence accumulation) parameter, we parsed a wide pool of human participants based on which planning strategy most likely accounted for their action selection strategy. Additional analyses corroborated the model classifications. Participants allocated to the route group (complex planning) were constrained by a higher decision criterion (boundary), and uniquely showed slower RT when planning segmental routes, consistent with the idea that they needed to incorporate more bits of information into their choices. This group also showed an enduring bias away from using the high-cost cursor.

After parsing participants on their likely decision strategies, we next investigated two core hypotheses. The first tested whether the efficiency of decision heuristics extends to action- execution contexts. Here, our findings confirmed that in a task requiring exquisite spatio-temporal control of selected actions, decision heuristics nonetheless required fewer trials to achieve better choice optimality and aligned with higher overall reward yields. Next, we probed how decision heuristics relate to individual differences in skill, measuring the latter in terms of independent dimensions of spatial and temporal precision. Here, we observed that heuristic adoption aligned with better spatial skill with high-cost actions. Together with the route group showing a parametric bias away from the high-cost cursor, we interpret the combined data across our task’s action- execution contexts as unambiguously supporting heuristic adoption in higher-skilled systems.

These combined findings extend the remit of heuristics to include contexts involving nontrivial action and further propose that they primarily preserve efficiency through less-is-more principles. Specifically, participants using the heuristic showed combined decisional and skill advantages that would not be predicted under accuracy-resource trade-off principles. A core rationale of sensorimotor-based forward models of action selection is that efference copies and predicted sensorimotor costs provide improved efficiency and robustness in the face of noisy sensory- prediction errors^27–29^. A corollary is that a low-skill system will struggle to accurately parameterise a complex action plan, increasing computational requirements during the planning phase and/or generating a high volume of online noise-driven corrective action during action execution^30, 31^. In either case, accuracy-resource trade-off principles would predict heuristic efficiency under a compensatory model, i.e., that the low-skill system is better served by adopting a simpler heuristic decision strategy that avoids fruitless deployment of computational resources.

Instead, under less-is-more principles, we propose that heuristic adoption in selection-execution contexts aligns with progress along motor-learning trajectories previously observed in forced-choice motor skill tasks (i.e., no selection required)^13, 14^. Here, early in the acquisition of novel motor skills, internal models that simulate action outcomes can expedite learning in exchange for high computational cost^13^. As participants then amass a wider cache of state transitions and successful experiences, control shifts from deliberative model-based planning to less taxing draws of state-appropriate motor outputs from memory^13^. Our findings are consistent with action selection following a similar qualitative trajectory across individuals; participants with superior skill, i.e., farther along motor-learning trajectories, also used a less taxing policy to select actions.

Under similar less-is-more principles we additionally propose that a cognitive substrate of heuristics might be agency. In computational terms, the core difference between our task and paradigms previously exploring heuristics is the source (internal vs external) of its generative model. Trial outcomes in our task were determined solely by a joint function that integrated participants’ cursor selection and its subsequent execution. In other words, outcome variance was fully determined by parameters (decision and performance) generated intrinsically by participants. In contrast, forecasting and computerised emulations typically employ extrinsic generative models, where outcome variance is a function of parameters beyond participants’ control. Recent evidence from bandit tasks (a computerised emulation with an extrinsic generative model) further suggests that humans might overparameterise their choices when extrinsic forces determine their fate, resulting in apparently irrational summary behaviour such as probability matching^32^. However, probability matching dissipates as a function of increased agency, for example, with increased motor involvement in choice execution^32^. While it is premature to conclude that increased agency will globally drive the adoption of heuristics, our findings nonetheless predict that a low-skill system (i.e., low agency) will more likely overparameterise choices in a selection- execution context, than revert to heuristics. This framework, pitting parameterisation against agency, is also consistent with emerging associations in clinical computational work, where sequelae such as overthinking (in anxiety^33^) and rumination (in depression^34^) align with excess deliberative model-based learning^35^.

We reveal additional evidence that planning dimensionality and skill might not simply evolve independently along separate strands of a learning manifold. In our task, we were able to probe how the adoption of heuristics or complex planning aligned with the dimensions shaping both skill state and skill learning. As mentioned above, in terms of skill state, we first observed that the heuristic group’s skill advantage was localised to the spatial dimension, with the groups not differing in terms of temporal skill. The heuristic group was therefore more skilled solely in the core (spatial) feature that characterised the decision policy best describing their action-selection data. In a series of time-on-task analyses (skill learning) we additionally observed that the route group, likely employing more complex planning, demonstrated skill learning across a broad array of motor-control features, including learning in temporal task dynamics. The heuristic group, in contrast, only showed skill learning in either the spatial or overall reward realms, i.e., no skill learning in the temporal domain (corroborated in a follow-up nonparametric analysis). Of note, a third nonplanner group, who never incorporated any state parameters into choice, and largely exploited the low-cost action, nonetheless improved across dimensions of motor-skill, including temporal skill (albeit only with the congruent cursor). In other words, the only group not showing credible plasticity in temporal skill learning was the heuristic group, i.e., the group who likely did not incorporate this information when selecting actions.

These combined findings support the idea that a yoked dimensionality might exist between policy governing the selection of actions and the skill shaping their subsequent execution, that is, the dimensions of information guiding action selection might be yoked to salient features in skill learning. In terms of a bottom-up framework, the spatial dominance of the heuristic group’s skill advantage, and likely spatial focus during action selection, suggests that such dimensional yoking may be modulated by skill-first credit assignment^36^. In other words, higher execution proficiency stemming predominantly from spatial precision may have overweighted this dimension during planning. Previous research has indeed shown that human choice policy can be separately influenced by distinct dimensions of error depending on the reliability of their signals^37–39^, and that increased agency might determine whether policy integrates either motor or reward-based errors^10^. However, an alternative top-down framework is also supported by the apparent absence of temporal learning in the heuristic group. Note that the route and heuristic groups did not differ in terms of overall temporal skill, just that the heuristic group uniquely showed no time-on-task evolution in this domain. An intriguing implication of this pattern of results is that a controller that localises a cardinal subset of information for making state-appropriate action selections might itself be able to influence controllers of what it considers superfluous features of sensorimotor error.

Future behavioural enquiry into heuristics could employ advancements in the selection-execution framework to investigate the yoked dimensionality hypothesis and investigate its potential bottom-up and top-down underpinnings in more detail. An additional key outstanding question relates to the robustness of heuristic adoption over time. Given the tendency for learning-related configurations in the human brain to vary more across rather than within individuals^40^, we employed a between-groups analytic approach inspired by an increasing body of work that uses behavioural profiles to cluster groups of individuals to increase robustness and reliability of hypothesis-specific brain activity^24, 41, 53^. While the present DDM parsimoniously distinguished human participants based on strategy-specific modulations of drift-rate, parameters were necessarily static. Our data therefore cannot inform any within-subject hypotheses regarding heuristic adoption; whether, for example, the route group would eventually reduce planning dimensionality with increased time-on-task. Though supplementary logistic models revealed the route group’s bias toward the low-cost cursor endured in later runs, suggesting their planning strategy may have held firm across the experiment, we cannot confirm whether they demonstrated a robust phenotypic trait or a relatively slower evolution along a trajectory of policy formation mutually traversed by both them and the heuristic group.

## Conclusion

The association between decision heuristics and intrinsic skill has historically evaded evaluation due to the simplified nature of action in computerised goal-oriented tasks. Here we used a novel task emulating both the decisional and skill-based demands of goal-oriented behaviour in a dynamic environment. The DDM parsimoniously identified human participants who likely adopted heuristics, and later modelling unambiguously aligned this lower-dimensional planning strategy with higher skill, consistent with less-is-more principles. We additionally observe that the intricacy of planning potentially maps onto the granularity of improvements in skill. Advancements in the behavioural assays of actions selected and executed will hopefully uncover the underlying causality, learning dynamics and neural underpinnings giving rise to this possible yoked dimensionality.

## Materials and Methods

### Participants and overview

We report pooled data from two experiment cohorts, with 53 right-handed human participants recruited in total, via both word-of-mouth and the online participant-recruitment portal at the University of California, Santa Barbara (UCSB). 34 participants reported as female and the group had an average (standard deviation) age of 21.9 (3.05) years. Participants performed the experiment either in a behavioural-testing suite (cohort 1, n=16) or an fMRI context (cohort 2, n=37). We report only behavioural data in the present paper from both groups. Visual angle subtended by stimuli was constant for the two cohorts and neither cohort differed in terms of eventual DDM group classifications (see Supplementary Materials: *Cohort-specific DDM group classifications*). Participant remuneration was $10 ($20, cohort 2) per hour baseline rate, with an additional $10 ($20, cohort 2) contingent on performance. Testing took place during a single session. The Institutional Review Board at UCSB approved all procedures (Human Subject Committee protocol: 36-21-0405). Prior to participating, participants provided informed written consent. All stimuli were presented using freely available functions^42, 43^ written in MATLAB code, and unless otherwise stated all analyses were also conducted using custom MATLAB scripts.

### Action selection-execution task: boatdock Paradigm

Our task was a continuous, nonlinear adaptation of the discrete grid-sail task^13^, extended such that reward yields require joint optimisation of action selection and action execution. All visual stimuli appear on a screen with a gray background (RGB_[0,1]_=[0.500,0.500,0.500]). In each trial (Figure 1a), they select one of two cursors, depicted by equilateral triangles (side length=0.830 °), to pilot from a start (S) to a goal (G), respectively depicted by a black (RGB_[0,1]_=[0,0,0]) and white (RGB_[0,1]_=[1,1,1]) square (side length=1.37 °). The SG pair appears within a circular grid (radius=3.82 °) centred on the screen centre. Locations of the SG are drawn with uniform probability on each trial, constrained such that neither element falls within 0.320 ° of the grid perimeter, and their centres are at least 0.957 ° apart.

Each cursor displaces in three deterministic directions (Figure 1b.), mapping onto the same three separate response buttons (“throttles”) operated by the right hand for the duration of the experiment (Figure 1c). One “congruent” cursor displaces at angles 7π/6 (index finger), π/2 (middle finger), and 11π/6 (ring finger) in a reference frame where π/2 aligns with the vertical meridian of the screen (Figure 1c). The other “incongruent” cursor displaces at angles 5π/6, π/6 and 3π/2, via one of two sets of spatially incongruent throttle-mappings, selected with uniform (p=0.500) probability for each subject (an example mapping is in Figure 1c). For the entire experiment, the congruent and incongruent cursors are identified by a different colour, green RGB_[0,1]_=[0,1,0] and blue RGB_[0,1]_=[0,0,1], determined with uniform (p=0.500) probability before each participant’s session.

For every frame a single throttle is down, the cursor will accelerate in that direction (see Supplementary Materials for specific acceleration dynamics) and a one unit of fuel is also subtracted from an allocation of 360 units provided for each trial. Participants therefore have a total 6 s throttle time on each trial before fuel depletes (refresh rate=60 Hz). Following a successful “dock” (see below) a screen informs participants of the fuel conserved, expressed as a proportion of the starting tank. No other exogenous cue is provided to participants regarding the size of the initial fuel allocation, or its rate of depletion.

### Trial structure

Each trial initiates with the action-selection period, signified by the appearance of an SG pair within a grid (“action selection”, Figure 1a). Participants have no time limit to select their desired cursor with the middle or index finger of their left hand, respectively using “a” or “z” of a standard keyboard (site 1) or buttons 1 and 2 (i.e., the two most leftward) of a six-button bimanual response box^44^ (site 2). Finger-cursor mapping (i.e., index→congruent, middle→incongruent, or vice versa) is determined every twenty trials by uniform (p=0.500) probability, prompted throughout the action-selection period by a silhouette of a hand (9.49 °-by-9.49 °) below the grid, with the relevant cursor above the relevant finger. Once an action is selected, the action-execution period immediately begins, signified by the silhouette prompt disappearing and the selected cursor spawning at the centre of S (“trial start”, Figure 1a). Participants now pilot the cursor from S to G with their right hand, using the “v” (index), “h” (middle) or “m” (ring) buttons on the keyboard (site 1) or buttons 4-6 on the right side of the response box (site 2). Action execution lasts until one of four possible trial outcomes. A successful “dock” is achieved if the cursor enters a 0.479 °- radius circular threshold (not visible to participants) centred on the centre of G, at a velocity no greater than 1.920 °/s. Alternatively, three catastrophic errors can occur if participants (i) run out of fuel, i.e., cumulative throttle time greater than 6 s; (ii) leave the grid; or (iii) enter the circular G threshold at a velocity greater than 1.920 °/s. Once a trial outcome is achieved, a feedback screen immediately informs participants of the outcome, respectively, “WELL DONE!”, “OUT OF GAS!”,“LEFT THE GRID!” or “TOO FAST!”, presented at the centre of the screen along with “SCORE: $”, where $ is either the proportion of fuel preserved (for successful docks) or 0 for all catastrophic errors. The feedback remains on the screen for 1 s, followed by a blank grey inter-trial-interval screen lasting one, two or three seconds (determined on each trial with uniform probability p=0.333). Participants performed 360 choice trials in total, portioned into six runs of 60 trials. Interlaced between choice runs were 20 practice trials, on which scores do not count toward the final bonus, forcing ten trials with both the congruent and incongruent cursor in pseudorandom order.

### Dependent variables

To enumerate cursor-state suitability a simple spatial heuristic we computed the angles (in °) between the vector of a trial’s SG and each vector on the incongruent cursor. The vector creating the smallest angle (which we term the “offset”) quantifies cursor suitability from this heuristic on a raw scale where values close to 0 reflect an SG perfectly aligning with one of the incongruent vectors and values close to 60 reflect an SG perfectly aligning with one of the congruent vectors. To enumerate action values derived from more complex planning we first computed forward simulations of the optimal routes on each (simulation procedure described in Supplementary materials). We subtract the total frames spent accelerating during the optimal route (λ) from the starting fuel bank of 360 units to estimate the maximum reward obtainable on a given trial.

We enumerated reaction time (RT) for action selection as the time elapsed between the time of the first frame of the action-selection screen (described above) and the time of cursor selection. We coded optimal selection as incongruent cursor on trials with offset<30 and congruent cursor on trials with offset>30 (optimal cursor selection did not differ depending on which action value (route vs heuristic) is computed).

We enumerated skill performance on each trial in terms of reward, spatial action execution and temporal action execution. Reward was the amount of fuel conserved. All modeling of reward used raw units (i.e., on a scale of 0 to 360) to allow Gaussian likelihood functions, however for clarity in reported results we present findings as a proportion of the tank preserved (from 0 to 1). Spatial action execution was the number of direction changes, i.e., a count of how many times a different throttle was pressed relative to the one previous. Temporal action execution was the difference between the cursor’s maximum velocity recorded during action-execution (in °/s), and the final velocity (in °/s) taken at the moment the cursor crossed the circular threshold around G, normalised by the distance covered by the SG (in °).

### Data analysis Computational modeling

We modeled action planning leading up to cursor selection with variants of a standard drift- diffusion model^45–47^. The full models included five free parameters: high-difficulty drift-rate *μ_1_*, low-difficulty drift-rate *μ_2_*, boundary *B*, congruency bias *b_C_*, and nondecision time *t_0_*. The boundaries for congruent and incongruent choices were defined as *B* and *-B*, respectively, and the starting point for the stochastic process was *b_C_B*. Parameters were necessarily constrained as follows: 0 ≤ *μ_1_* ≤ *μ_2_*, *μ_2_* ≥ 0, *B* > 0, -1 < *b_C_* < 1, and *t_0_* ≥ 0. Noise was represented as the standard deviation of diffusion with a fixed scaling parameter *σ* = 0.1.

We compared three types of models: two route-planning models (with one or two drift-rates), two heuristic models (with one or two drift-rates), and the null (i.e., nonplanning) model. For route-planning models, we determined difficulty by dividing trialwise differences in reward yields (between the simulated optimal routes for either cursor) into five bins. For the heuristic models, we determined difficulty by dividing trialwise offsets into five bins. The five difficulty bins corresponded to drift-rates of -*μ_2_*, -*μ_1_*, 0, *μ_1_*, and *μ_2_*. We constrained single-drift-rate models such that *μ_1_*= *μ_2_* to minimise penalties for additional degrees of freedom, and the null model such that *μ_1_*= *μ_2_* = 0 to represent insensitivity to the onscreen information.

We fitted candidate models to empirical distributions of choices and RTs at the level of individual subjects using maximum-likelihood estimation and the “chi-square” fitting method^48^. We calculated the frequencies of either choice and the 10, 30, 50, 70, and 90% quantiles (i.e., six bins) of their respective RT distributions for each difficulty level. Free parameters were optimised with respect to overall goodness of fit for a given subject using iterations of the Nelder-Mead simplex algorithm with randomised seeding^49^. We adjusted for model complexity when comparing models that differed in degrees of freedom using the Akaike information criterion with correction for finite sample size (AICc)^25, 26^.

Three participant groups were defined by the results of model fitting following penalisation. The “heuristic” and “route” groups included those who were best fitted by a heuristic or route-planning model, respectively, according to the AICc. Assignment to the “nonplanner” group meant that adding free parameters for planning did not yield a significant improvement in goodness of fit relative to the null model containing no sensitivity to either heuristic or route-based enumeration of cursor suitability on each SG.

### Bayesian models

We sampled all Bayesian posterior distributions using No U-Turn sampling (NUTS) Hamiltonian Monte Carlo, implemented with the PyMC3 package^50^ in custom Python scripts. Unless otherwise specified, each model’s posterior distributions were sampled across four chains of 10000 samples (40000 total), with an additional initial 10000 samples per chain (40000 total) discarded after tuning the sampler’s step-size to an acceptance threshold of 0.95 (80000 samples combined), with further convergence criteria that no chains contain any divergences and no posterior’s *R̂* value, estimating the ratio of variance within the n=4 chains to the variance of the pooled chains, greater than 1 (see:^51^). Unless otherwise stated, dependent variables were z-score normalised across participants prior to fits. We calculated minimum-width Bayesian credible intervals of relevant posteriors from their chains, using the default settings for Highest Density Interval (HDI) calculation in the arviz package^52^.

We fitted summary models to variables that could first be summarised across trials. The model for median RT assumed individual participant (n) values (y) were characterised by a Gaussian likelihood function, i.e., *y*_*n*_ ∼ Ɲ(μ, Σ). Median RT variables were z-score normalised across all subjects prior to fitting, and we respectively assigned μ and Σ an uninformed Gaussian and half-Gaussian prior: μ ∼Ɲ(0,10) and Σ∼halfƝ(10). We report the expected value (𝔼(x)) and highest density interval (HDI(x)) of μ, i.e., group-mean RT.

Three separate binomial models then estimated summaries of behaviour as measured by three binomial variables p(congruent cursor), p(optimal cursor) and p(catastrophic error). Each of these summary models used a Binomial likelihood function y∼Bin(θ,t), where y and t are n-element vectors, respectively enumerating the number of observed instances reported by each individual participant (n) and their total number of trials (t). In each model, we assigned θ an uninformed prior from the beta distribution: θ ∼ Beta(α=1,β=1). In each case we report the expected value (𝔼(x)) and highest density interval (HDI(x)) of θ, i.e., respectively, group-mean p(congruent cursor), p(optimal cursor) and p(catastrophic error).

After using the DDM to classify participants into groups (heuristic, route, nonplanner) we then ran modified versions of the above models, to compute group-specific posteriors (reported in Table 1). The median RT model assumed individual participant (n) values (y) for each variable (v) was characterised by a separate Gaussian likelihood function, further depending on n’s group-allocation (g(n): heuristic, route, nonplanner), i.e., *y*_*n*_ ∼ Ɲ(μ *_g(n)_*,Σ*_g(n)_*). Each variable was z-score normalised separately (but across all subjects) prior to fitting, and we respectively assigned each μ*_,g(n)_* and Σ*_g(n)_* an uninformed Gaussian and half-Gaussian prior: μ *_g(n)_* ∼Ɲ(0,10) and Σ*_g(n)_* ∼halfƝ(10). We report the expected value (𝔼(x)) and highest density interval (HDI(x)) of μ *_g(n)_*, i.e., group-specific median RT.

Three separate binomial models then estimated summaries of behaviour as measured by three binomial variables: p(congruent cursor), p(optimal cursor) and p(catastrophic error). For the n(g) participants in each group (g), each summary model used a Binomial likelihood function y_g_∼Bin(θ_g_,t_g_), where y_g_ and t_g_ are n(g)-element vectors, respectively enumerating the number of observed instances reported by each individual participant in a group (y_g_) and their total number of trials (t_g_). In each model, we assigned each θ_g_ an uninformed prior from the beta distribution: θ_g_∼ Beta(α=1,β=1). In each case we report the expected value (𝔼(x)) and highest density interval (HDI(x)) of θ_g_, i.e., respectively, group-specific p(congruent cursor), p(optimal cursor) and p(catastrophic error).

We then estimated group-specific DDM parameters (reported in Table 1). A single Gaussian model summarised seven variables in total. First, the three variables applicable to each group identified by the DDM framework, specifically: boundary - *B*, congruency bias - *b_C_*, and nondecision time - *t_0_*. In addition, the three variables applicable only to the route and heuristic groups, specifically: drift-rate, hi difficulty - *μ_1_*, drift-rate, lo difficulty- *μ_2_* and sensitivity - *S*. This model assumed individual participant (n) values (y) for each variable (v) were characterised by a separate Gaussian likelihood function, further depending on n’s group-allocation (g(n): route, heuristic or nonplanner), i.e., *y*_*n*_∼Ɲ(μ *_v,g(n)_*,Σ*_v,g(n)_*). Each variable was z-score normalised separately (but across all subjects) prior to fitting, and we respectively assigned each Μ*_v,g(n)_*and Σ*_v,g(n)_* an uninformed Gaussian and half-Gaussian prior: μ *_v,g(n)_* ∼Ɲ(0,10) and Σ*_v,g(n)_* ∼halfƝ(10). In each case, we report the expected value (𝔼(x)) and highest density interval (HDI(x)) of μ *_v,g(n_*, i.e., group-specific value of each DDM parameter.

We next fitted hierarchical models that imposed hierarchical structures to summarise trialwise measures within each participant with participant-level posteriors that were themselves constrained by a relevant hierarchical parameter. We first used a hierarchical Bayesian binomial model to estimate the credible ranges of group-specific choice optimality (p(optimal cursor)), separately for each run. The hierarchical structure used Binomial likelihood functions to summarise the number of optimal cursor selections (y) made by each participant (n) for all trials (t) in a given run (r), y_n,r_∼Bin(θ_n,r_,t_n,r_). The model constrained θ_*n*,*r*_ posteriors with separate hierarchical group (g(n)) and run-specific Beta distributions, i.e.: θ_*n*,*r*_∼ Beta(α*_g(n),r_*,β*_g(n),r_*). Each α*_g(n),r_* and β*_g(n),r_*were assigned uninformed priors from a half-Student’s T distribution, i.e.: α*_g(n),r_* ∼ HalfStudentT(10,10) and β*_g(n),r_* ∼ HalfStudentT(10,10), bounded to never draw values of α*_g(n),r_*=0 or β*_g(n),r_*=0. Run-specific group-level deterministic posterior estimates of optimal choice (θ_g(*n*),*r*_) were calculated by drawing 10,000 independent samples (k) from relevant α*_g(n),r_* and β*_g(n),r_* posteriors and computing the mean of the resulting kth Beta distribution, i.e., θ_g(*n*),*r*,k_ = α*_g(n),r,k_* / (α*_g(n),r,k_ +*β*_g(n),r,k_*). We report the expected value (𝔼(x)) and highest density interval (HDI(x)) of θ_g(*n*),*r*_, i.e., group-by-run-specific p(optimal cursor).

We then used two separate hierarchical Bayesian Gaussian models to estimate the credible ranges of group-mean performance in the two continuous action-execution variables (reward and temporal skill), separately for each run, and separately again for each cursor. In each model, the hierarchical structure used Gaussian likelihood functions to summarise each (n) participant’s trialwise measures across all trials in a given run (r), separately for each cursor (c), i.e.: *y*_*n*,*r*,*c*_∼Ɲ(*x̅*_*n,r,c*_,exp(σ_*n,r,c*_)). The model constrained *x̅*_*n,r,c*_ and σ_*n,r,c*_ posteriors with separate hierarchical group (g(n)), run (r) and choice-specific (c) Gaussian distributions, i.e.: *x̅*_*n,r,c*_∼ Ɲ(μ*_g(n),r,c_*,Σ*_g(n),r,c_*) and σ_*n,r,c*_ ∼ Ɲ(μ*_g(n),r,c_*, *ψ*_*g(n),r,c*_). Each μ*_g(n),r,c_* and μ*_g(n),r,c_*were assigned uninformed Gaussian priors (∼Ɲ(0,10)), while each Σ*_g(n),r,c_* and *ψ*_*g(n),r,c*_ were assigned uninformed half-Gaussian priors (∼halfƝ(10)). We report the expected value (𝔼(x)) and highest density interval (HDI(x)) of μ*_g(n),r,c_*, i.e., group-by-run-by-choice-specific reward yields. Note that the model for reward was fitted to a continuous measure, scoring fuel conserved on a scale of 0 to 360, but for clarity, we adjusted runwise and collapsed HDIs (division by 360), also prior to computing any HDIs related to between-comparisons, to express results as a proportion of fuel preserved. Time-on-task betas, however, relate to unadjusted posteriors.

We used a hierarchical Bayesian Poisson model to estimate the credible ranges of group-mean performance in spatial skill, separately for each run, and separately again for each cursor. In each model, the hierarchical structure used Poisson likelihood functions to summarise each (n) participant’s trialwise direction changes across all trials in a given run (r), separately for each cursor (c), i.e.: *y*_*n*,*r*,*c*_∼Pois(exp(*x̅*_*n,r,c*_)). The model constrained *x̅*_*n,r,c*_posteriors with separate hierarchical group (g(n)), run (r) and cursor-specific (c) Gaussian distributions, i.e.: *x̅*_*n,r,c*_∼Ɲ(μ*_g(n),r,c_*,Σ*_g(n),r,c_*). μ*_g(n),r,c_* and Σ*_g(n),r,c_*were respectively assigned uninformed Gaussian (∼Ɲ(0,10)) and half-Gaussian priors (∼halfƝ(10)). We report the expected value (𝔼(x)) and highest density interval (HDI(x)) of μ*_g(n),r,c_*, i.e., group-by-run-by-choice-specific direction changes. For clarity in reported results, we re-adjusted runwise and collapsed HDIs (exponential transform), also prior to computing any HDIs related to between-comparisons, to discount the use of exp(*x̅*_*n,r,c*_) in the likelihood function. Time-on-task betas, however, relate to unadjusted posteriors.

For both the hierarchical Gaussian and Poisson skill models, separately for each group (g) and choice (c), we enumerated deterministic posteriors of overall skill level by averaging each posterior sample across runs, i.e., for each posterior sample, 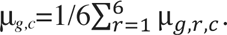. We then enumerated deterministic linear and logarithmic time-on-task effects b_g,c_ by drawing posterior samples from μ*_g,r,c_*. Specifically, on each (k) of 40,000 draws, we computed the kth column of b_g,c_ (b_g,c,k_), where b_g,c,k_=(X^T^X)^-1^X^T^Y_g,c,k_. Here, Y_g,c,k_ is a six-element column vector containing an independent draw from each run (r) of μ*_g,r,c_*and matrix X is a three-column matrix respectively containing six constant terms (1), z-scored linear x∈(1,2,…,6) and z-scored logarithmic x∈(ln(1),ln(2),…,ln(6)) regressors. The second and third rows of resulting 3-by-40,000 matrix b_g,c_ respectively contained deterministic posteriors for linear and logarithmic time-on-task effects. Where logarithmic time-on-task effects were credible (0 ∉ HDI), we considered that group-by- cursor time-on-task effect to be logarithmic even if a linear effect was also observed. Note, as specified above, that in reported results, we present the HDIs of time-on-task coefficients (linear and logarithmic) fitted to unadjusted runwise posteriors, i.e., before we made any adjustment to posteriors for intuitive presentation of runwise/collapsed HDIs.

For the individual-participant-level nonparametric analysis of temporal skill, we computed the median of each *x̅*_*n,r,c*_posterior from the relevant Gaussian skill model. Separately for each cursor we regressed the six-element vector of participant’s run-specific median values, first as a function of an intercept and a linear time-on-task regressor (z-scored linear x∈(1,2,…,6)), and then as a function of an intercept and a z-scored logarithmic regressor x∈(ln(1),ln(2),…,ln(6)). If either model’s regressor (x) was significant (determined by 95% coefficient confidence intervals not containing 0), we considered that participant time-on-task^+^ for that cursor and skill variable. We compared proportions of time-on-task^+^ participants (y) between groups (g), separately for each cursor, by fitting Binomial likelihood function y_g_∼Binomial(θ_g_,n_g_), assigning each θ_g_ an uninformed prior from the beta distribution: θ_g_∼ Beta(α=1,β=1).

In all above cases, we consider strong evidence of credible effects as follows: for comparison of parameters to criterion values (e.g., a regression coefficient above 0, or a likelihood above 0.50, etc.) we required the entire HDI of that parameter to not include the criterion value. For comparison of two parameters we required the HDI of the deterministic distribution of their difference (posterior A - posterior B) to not contain 0. Note that two HDIs might overlap, but that this deterministic distribution of difference may yet still not contain 0.

### Verifying DDM classification of heuristic and route groups with segmentation interaction

We tested whether the heuristic and complex planning elicited similar RT for SGs where the optimal solution was a single linear route from the start to the goal (single segment), but different RT where the optimal solution requires some kind of turn or zig-zag (multiple segment), i.e., where complex-planning demands likely become disproportionately greater than heuristic demands. On all trials where the optimal selection was the congruent cursor (i.e., offset>30 ° or route-planning score>0), we computed median RT for each participant separately for trials where our simulations (see Supplementary Materials: *Optimal route simulations*) returned a single-segment solution and for trials with a multiple-segment solution (see Figure 2i). We submitted this to a two-way ANOVA with factors group (heuristic, route) and segmentation (single, multiple) with significant effects probed using post-hoc tests applying Tukey correction. We also repeated this analysis for trials where the optimal selection was the incongruent cursor.

## Acknowledgements

The research was supported by award #W911NF-16-1-0474 from the Army Research Office and by the Institute for Collaborative Biotechnologies under Cooperative Agreement W911NF-19-2- 0026 with the Army Research Office.

## Competing interests

Authors declare no financial, and no nonfinancial, competing interests.

## Notes

### Competing Interest Statement

The authors have declared no competing interest.

### Summary of Updates

Rewrite to explain concepts with greater clarity

