## Supplementary Materials for "Decision heuristics in contexts exploiting intrinsic skill"

*Nonplanner group*

In addition to the 33 participants identified by our DDM framework as likely employing one of two goal-oriented strategies, 20 participants were best fitted by the null model (skill variables of this group are included in Supplementary Table 1). While our hypotheses principally focused on decisional and skill differences between the route and heuristic groups, we briefly comment here on the nonplanner group. Of all groups, nonplanners showed the fastest overall decision time and strongest DDM bias toward the congruent cursor (Table 1), consistent with an action-selection policy that did not integrate the external state. However, despite demonstrating no evidence of state-appropriate action selection (Figure 3b.), largely stemming from an over-reliance on the congruent cursor (Table 1), nonplanners nonetheless exhibited skill learning during the execution portion of our task (Figure 3b-d). They improved with both cursors in terms of reward yield and spatial precision, but only demonstrated improved temporal dynamics with the congruent cursor, i.e., the cursor they exploited to yield reward.

**Supplementary Table 1:** choice and skill measures; group-by-run, collapsed across runs, and time-on-task effects

| cursor/variable | posterior | heuristic | | route | | nonplanner | |
|---|---|---|---|---|---|---|---|
| | | $\mathbb{E}(x)$ | HDI(x) | $\mathbb{E}(x)$ | HDI(x) | $\mathbb{E}(x)$ | HDI(x) |
| p(optimal choice) | $\theta_{run1}$ | 0.596 | [0.366,0.823] | 0.555 | [0.354,0.760] | 0.472 | [0.279,0.658] |
| | $\theta_{run2}$ | 0.698 | [0.501,0.898]‡ | 0.638 | [0.433,0.830] | 0.493 | [0.307,0.681] |
| | $\theta_{run3}$ | 0.710 | [0.514,0.893]‡ | 0.668 | [0.469,0.848] | 0.493 | [0.306,0.689] |
| | $\theta_{run4}$ | 0.735 | [0.543,0.906]‡ | 0.712 | [0.533,0.882]‡ | 0.484 | [0.299,0.667] |
| | $\theta_{run5}$ | 0.740 | [0.557,0.908]‡ | 0.729 | [0.555,0.887]‡ | 0.484 | [0.297,0.668] |
| | $\theta_{run6}$ | 0.747 | [0.560,0.938]‡ | 0.718 | [0.536,0.885]‡ | 0.508 | [0.321,0.696] |
| cong/reward | $\mu_{run1}$ | 0.481 | [0.430,0.532] | 0.489 | [0.463,0.515] | 0.441 | [0.415,0.468] |
| | $\mu_{run2}$ | 0.531 | [0.484,0.580] | 0.530 | [0.504,0.557] | 0.466 | [0.442,0.488] |
| | $\mu_{run3}$ | 0.574 | [0.535,0.613] | 0.569 | [0.535,0.603] | 0.517 | [0.495,0.538] |

| | | | | | | | |
|---|---|---|---|---|---|---|---|
| | $\mu_{run4}$ | 0.571 | [0.533,0.608] | 0.567 | [0.528,0.608] | 0.507 | [0.484,0.530] |
| | $\mu_{run5}$ | 0.568 | [0.537,0.598] | 0.581 | [0.550,0.614] | 0.516 | [0.496,0.535] |
| | $\mu_{run6}$ | 0.577 | [0.543,0.610] | 0.590 | [0.562,0.619] | 0.529 | [0.504,0.554] |
| | $\mu_{all\ runs}$ | 0.550 | [0.533,0.566] | 0.554 | [0.542,0.567] | 0.496 | [0.487,0.505] |
| | $\beta_{lin}$ | 0.144 | [0.058,0.230]* | 0.16 | [0.103,0.218]* | 0.141 | [0.091,0.191]* |
| | $\beta_{log}$ | 0.36 | [-0.004,0.714] | 0.241 | [-0.003,0.480] | 0.205 | [0.004,0.397]* |
| incong/reward | $\mu_{run1}$ | 0.479 | [0.425,0.537] | 0.409 | [0.369,0.449] | 0.374 | [0.338,0.410] |
| | $\mu_{run2}$ | 0.516 | [0.461,0.568] | 0.491 | [0.450,0.532] | 0.416 | [0.375,0.462] |
| | $\mu_{run3}$ | 0.582 | [0.549,0.615] | 0.537 | [0.491,0.584] | 0.453 | [0.412,0.493] |
| | $\mu_{run4}$ | 0.569 | [0.528,0.608] | 0.558 | [0.519,0.584] | 0.478 | [0.446,0.510] |
| | $\mu_{run5}$ | 0.567 | [0.534,0.600] | 0.559 | [0.519,0.598] | 0.47 | [0.439,0.499] |
| | $\mu_{run6}$ | 0.572 | [0.536,0.690] | 0.549 | [0.515,0.601] | 0.486 | [0.437,0.533] |
| | $\mu_{all\ runs}$ | 0.548 | [0.530,0.565] | 0.517 | [0.500,0.534] | 0.446 | [0.431,0.462] |
| | $\beta_{lin}$ | 0.147 | [0.055,0.243]* | 0.225 | [0.146,0.308]* | 0.182 | [0.101,0.268]* |
| | $\beta_{log}$ | 0.379 | [0.008,0.763]* | 0.564 | [0.236,0.897]* | 0.267 | [-0.042,0.599] |
| cong/spatial | $\mu_{run1}$ | 2.13 | [1.70,2.65] | 2.04 | [1.82,2.28] | 2.76 | [2.50,3.04] |
| | $\mu_{run2}$ | 1.60 | [1.28,2.00] | 1.70 | [1.47,1.96] | 2.41 | [2.24,2.61] |
| | $\mu_{run3}$ | 1.43 | [1.15,1.77] | 1.49 | [1.27,1.73] | 2.07 | [1.89,2.26] |
| | $\mu_{run4}$ | 1.33 | [1.09,1.61] | 1.42 | [1.22,1.66] | 2.13 | [1.94,2.34] |
| | $\mu_{run5}$ | 1.41 | [1.17,1.69] | 1.34 | [1.18,1.52] | 2.02 | [1.84,2.21] |
| | $\mu_{run6}$ | 1.38 | [1.15,1.64] | 1.24 | [1.08,1.43] | 1.89 | [1.69,2.10] |
| | $\mu_{all\ runs}$ | 1.53 | [1.41,1.66] | 1.52 | [1.43,1.61] | 2.20 | [2.11,2.28] |
| | $\beta_{lin}$ | -0.128 | [-0.212,-0.048]* | -0.157 | [-0.211,-0.105]* | -0.117 | [-0.156,-0.076]* |
| | $\beta_{log}$ | -0.369 | [-0.703,-0.024]* | -0.16 | [-0.378,0.054] | -0.145 | [-0.303,0.007] |
| incong/spatial | $\mu_{run1}$ | 2.13 | [1.702,2.686] | 2.79 | [2.438,3.190] | 3.38 | [3.053,3.736] |
| | $\mu_{run2}$ | 1.68 | [1.324,2.106] | 1.95 | [1.650,2.323] | 2.83 | [2.354,3.380] |
| | $\mu_{run3}$ | 1.39 | [1.143,1.696] | 1.63 | [1.384,1.916] | 2.38 | [2.018,2.782] |
| | $\mu_{run4}$ | 1.45 | [1.174,1.766] | 1.47 | [1.265,1.706] | 2.42 | [2.123,2.748] |
| | $\mu_{run5}$ | 1.45 | [1.220,1.726] | 1.48 | [1.287,1.713] | 2.44 | [2.130,2.804] |
| | $\mu_{run6}$ | 1.51 | [1.204,1.893] | 1.57 | [1.380,1.791] | 2.23 | [1.895,2.654] |
| | $\mu_{all\ runs}$ | 1.58 | [1.452,1.726] | 1.76 | [1.662,1.874] | 2.59 | [2.439,2.748] |
| | $\beta_{lin}$ | -0.104 | [-0.195,-0.015]* | -0.185 | [-0.241,-0.128]* | -0.121 | [-0.180,-0.063]* |
| | $\beta_{log}$ | -0.392 | [-0.744,-0.035]* | -0.499 | [-0.728,-0.262]* | -0.228 | [-0.458,-0.008]* |
| cong/temporal | $\mu_{run1}$ | 0.655 | [0.598,0.712] | 0.633 | [0.579,0.688] | 0.694 | [0.650,0.736] |

| | | | | | | | |
|---|---|---|---|---|---|---|---|
| | $\mu_{run2}$ | 0.627 | [0.555,0.700] | 0.661 | [0.609,0.711] | 0.654 | [0.619,0.692] |
| | $\mu_{run3}$ | 0.649 | [0.571,0.733] | 0.683 | [0.628,0.738] | 0.680 | [0.640,0.721] |
| | $\mu_{run4}$ | 0.686 | [0.610,0.761] | 0.694 | [0.643,0.743] | 0.712 | [0.674,0.749] |
| | $\mu_{run5}$ | 0.705 | [0.637,0.776] | 0.744 | [0.659,0.793] | 0.723 | [0.688,0.757] |
| | $\mu_{run6}$ | 0.701 | [0.621,0.784] | 0.724 | [0.665,0.787] | 0.739 | [0.710,0.768] |
| | $\mu_{all\ runs}$ | 0.671 | [0.641,0.700] | 0.690 | [0.667,0.712] | 0.700 | [0.685,0.716] |
| | $\beta_{lin}$ | 0.086 | [-0.018,0.187] | 0.124 | [0.043 0.205]* | 0.080 | [0.027 0.133]* |
| | $\beta_{log}$ | -0.124 | [-0.530,0.279] | 0.062 | [-0.250,0.385] | -0.184 | [-0.411,0.034] |
| incong/temporal | $\mu_{run1}$ | 0.719 | [0.638,0.798] | 0.621 | [0.565,0.676] | 0.722 | [0.623,0.819] |
| | $\mu_{run2}$ | 0.713 | [0.647,0.780] | 0.664 | [0.612,0.713] | 0.721 | [0.660,0.787] |
| | $\mu_{run3}$ | 0.684 | [0.612,0.750] | 0.651 | [0.601,0.701] | 0.693 | [0.630,0.757] |
| | $\mu_{run4}$ | 0.680 | [0.611,0.751] | 0.675 | [0.612,0.741] | 0.767 | [0.686,0.848] |
| | $\mu_{run5}$ | 0.727 | [0.638,0.811] | 0.698 | [0.630,0.763] | 0.768 | [0.697,0.835] |
| | $\mu_{run6}$ | 0.723 | [0.660,0.783] | 0.751 | [0.689,0.814] | 0.805 | [0.750,0.862] |
| | $\mu_{all\ runs}$ | 0.708 | [0.678,0.737] | 0.677 | [0.654,0.701] | 0.746 | [0.716,0.776] |
| | $\beta_{lin}$ | 0.010 | [-0.096,0.114] | 0.134 | [0.048,0.217]* | 0.110 | [-0.004,0.219] |
| | $\beta_{log}$ | -0.181 | [-0.592,0.232] | -0.074 | [-0.389,0.256] | -0.208 | [-0.657,0.270] |

*Notes: non=nonplanner; heur=heuristic; cong=congruent cursor; incong=incongruent cursor; spatial=spatial skill; temporal=temporal skill; coll.=collapsed across runs; runwise and collapsed HDIs for reward have been re-adjusted (division by 360) to express reward as a proportion of fuel preserved; *=time-on-task coefficient credibly dearts 0;+ proportion of choices credibly above chance optimality (0.50).*

*Hierarchical logistic choice model*

We used a hierarchical Bayesian logistic regression model to assess the group-specific modulation of choice (p(incongruent)) as a function of an intercept (β0), trial offsets (β1; i.e., the trialwise enumeration of heuristic value) and the Euclidean distance (in screen pixels) of trial SGs (β2). The hierarchical structure used Bernoulli likelihood functions to characterise choice likelihood for each individual participant (n) and trial (t), i.e.: $y_{n,t}$~Bernoulli($p_{n,t}$), where $p_{n,t}$ is computed with a deterministic logistic transition function $S(x_{n,t})$, where $x_{n,t}=\beta0_n+\beta1_n*\text{offset}_{n,t}+\beta2_n*\text{distance}_{n,t}$. The model constrained coefficient posteriors fitted to each participant's set of trials with separate hierarchical group-specific (g(n)) Gaussian distributions, i.e.: $b0_n \sim \mathcal{N}(\beta0_{g(n)},\Sigma_{0,g(n)})$, $b1_n \sim \mathcal{N}(\beta1_{g(n)},\Sigma_{1g(n)})$ and $b2_n \sim \mathcal{N}(\beta2_{g(n)},\Sigma_{2g(n)})$. Each $\beta0_{g(n)}$, $\beta1_{g(n)}$ and $\beta2_{g(n)}$ were assigned uninformed Gaussian priors ($\sim\mathcal{N}(0, 10)$), while each $\Sigma_{0g(n)}$, $\Sigma_{1g(n)}$ and $\Sigma_{2g(n)}$ were assigned uninformed half-Gaussian priors (~half$\mathcal{N}(10)$). Both regressors were z-score normalised across all trials from all subjects prior to fitting. Finally, we fitted two iterations of this model, one using trials from the early phase of the task (first three runs), and a second using trials from the late phase of the task (final three runs).

Results of this hierarchical logistic regression model are summarised below in Supplementary Table 2. This model first bolstered the DDM by demonstrating the route and heuristic group uniquely integrated state information into action selection. During both early and late phases of the task, the route (HDI($\beta1_{\text{route,early}}$)=[-0.865,-0.483]; HDI($\beta1_{\text{route,late}}$)=[-1.559,-1.014]) and heuristic group (HDI($\beta1_{\text{heuristic,early}}$)=[-1.466,-0.601]; HDI($\beta1_{\text{heuristic,late}}$)=[-2.080,-1.201]), incorporated route offsets optimally into choice; note that their credibly negative coefficient HDIs reflect increased likelihood of selecting the incongruent cursor when offset angle was low, i.e., suited to the incongruent cursor (offset was normalised to vectors on the incongruent cursor; see: Methods/Results). In addition, this model supported the finding from the DDM relating to the route group's bias. The route group uniquely showed a bias to the congruent cursor in both early and late phases of the task, (HDI($\beta0_{\text{route,early}}$)=[-0.682,-0.240]; HDI($\beta0_{\text{route,late}}$)=[-0.423,-0.104]), which was not credibly evident in the heuristic group in either instance (HDI($\beta0_{\text{heuristic,early}}$)=[-0.480,0.011];

HDI($\beta0_{heuristic,late}$)=[-0.401,0.073]). No groups credibly modulated their choice by the distance covered by a route's start-goal pairing (SGSG), in either early or late phases of the task (all HDIs for β2 subtend 0 in Supplementary Table 2). However, of note, the trending positive distance parameter estimate for the route group in the early phase (HDI($\beta2_{route,early}$)=[-0.015,0.136]) suggests first that their planning strategy may not have been born out of risk-aversion, (which instead would have been characterised by incongruent selection on shorter SGs). Though we can only speculate on a non-credible finding, if the route group selectively used the high-cost incongruent cursor early in primarily longer SGs, they may have been reserving its usage for situations where optimal choice was disproportionately beneficial, due to the nonlinear task physics.

**Supplementary Table 2:** Hierarchical logistic model of choice behaviour parameters

| task phase | parameter | heuristic | route | nonplanner |
|---|---|---|---|---|
| | | HDI(x) | HDI(x) | HDI(x) |
| early | β0 | [-0.480,0.011] | [-0.682,-0.240]* | [-1.690,-0.780]* |
| late | β0 | [-0.401,0.073] | [-0.423,-0.104]* | [-1.471,-0.672]* |
| early | $\beta1_{offset}$ | [-1.466,-0.601]* | [-0.865,-0.483]* | [0.016,0.196]* |
| late | $\beta1_{offset}$ | [-2.080,-1.201]* | [-1.559,-1.014]* | [-0.059,0.172] |
| early | $\beta2_{distance}$ | [-0.175,0.045] | [-0.015,0.136] | [-0.092,0.100] |
| late | $\beta2_{distance}$ | [-0.112,0.123] | [-0.083,0.087] | [-0.127,0.071] |

*Notes: non=nonplanner; heur=heuristic; int.=intercept; *=coefficient credibly departs 0;*

*Hierarchical Poisson model with choice-normalised spatial skill*

To dissociate the heuristic group's superior spatial skill with the incongruent cursor from their overall more optimal choice behaviour, we used a hierarchical Bayesian Poisson model to estimate the credible ranges of group-mean performance in spatial skill, using a measure which had been normalised by the optimal number of direction changes in the simulated solution (see Supplementary Materials: *Optimal route simulations*). This normalisation took the number of direction changes made on each trial, and subtracted from that the number of direction changes made by the optimal solution for the specific cursor chosen on that trial (i.e., not necessarily normalised to the optimal cursor for a given route, but the selected cursor). Due to a small number of resulting trials (0.3%, across all subjects) containing a negative value (never lower than -1), we added a constant (1) to all trials, to ensure the lowest value was 0, suitable for a Poisson likelihood function. With this normalisation, higher values reflect worse spatial skill, i.e., more direction changes relative to cursor-optimal. As with the unnormalised model, we fitted the model separately for each run, and separately again for each cursor. In each model, the hierarchical structure used Poisson likelihood functions to summarise each (n) participant's trialwise direction changes across all trials in a given run (r), separately for each cursor (c), i.e.: $y_{n,r,c} \sim \text{Pois}(\exp(\mu_{n,r,c}))$. The model constrained $\mu_{n,r,c}$ posteriors with separate hierarchical group (g(n)), run (r) and cursor-specific (c) Gaussian distributions, i.e.: $\mu_{n,r,c} \sim \mathcal{N}(M(\mu)_{g(n),r,c}, \Sigma(\mu)_{g(n),r,c})$. $M(\mu)_{g(n),r,c}$ and $\Sigma(\mu)_{g(n),r,c}$ were respectively assigned uninformed Gaussian ($\sim\mathcal{N}(\mu=0,\sigma=10)$) and half-Gaussian priors ($\sim\text{half}\mathcal{N}(\sigma=10)$). For clarity in reported results, we re-adjusted runwise and collapsed HDIs (exponential transform, followed by subtraction of -1), also prior to computing any HDIs related to between-comparisons, to discount first the use of $\exp(\mu_{n,r,c})$ in the likelihood function, and then the constant added to all trials prior to fitting. Time-on-task betas, however, relate to unadjusted posteriors.

Results of this model are summarised below in Supplementary Table 3. Crucially, collapsing across runs, we see the heuristic group demonstrating credibly fewer direction changes with the incongruent cursor ($\mathbb{E}(\mu_{heuristic-route})$=-0.174, $HDI(\mu_{heuristic-route})$=[-0.331,-0.005]), supporting the interpretation of the finding from the main paper (see: Results - *Comparisons of skill between route and heuristic groups*) that their superior incongruent spatial skill is independent to the navigational consequences of their choices.

**Supplementary Table 3:** spatial skill, normalised by cursor selection; group-by-run, collapsed across runs, and time-on-task effects

| cursor/skill | posterior | heuristic | | route | | non | |
|---|---|---|---|---|---|---|---|
| | | $\mathbb{E}(x)$ | HDI(x) | $\mathbb{E}(x)$ | HDI(x) | $\mathbb{E}(x)$ | HDI(x) |
| cong/spatial | $\mu_{run1}$ | 1.4 | [1.000,1.886] | 1.29 | [1.085,1.522] | 1.92 | [1.664,2.216] |
| | $\mu_{run2}$ | 0.9 | [0.610,1.228] | 0.96 | [0.742,1.195] | 1.59 | [1.408,1.776] |
| | $\mu_{run3}$ | 0.75 | [0.481,1.046] | 0.75 | [0.555,0.972] | 1.25 | [1.092,1.430] |
| | $\mu_{run4}$ | 0.63 | [0.422,0.872] | 0.7 | [0.519,0.891] | 1.26 | [1.073,1.479] |
| | $\mu_{run5}$ | 0.68 | [0.477,0.889] | 0.6 | [0.464,0.758] | 1.15 | [0.982,1.323] |
| | $\mu_{run6}$ | 0.63 | [0.432,0.861] | 0.51 | [0.363,0.669] | 1.06 | [0.879,1.261] |
| | $\mu_{all\ runs}$ | 0.814 | [0.704,0.926] | 0.784 | [0.707,0.861] | 1.35 | [1.276,1.436] |
| | $\beta_{lin}$ | -0.115 | [-0.178,-0.052]* | -0.132 | [-0.173,-0.092]* | -0.112 | [-0.148,-0.076]* |
| | $\beta_{log}$ | -0.274 | [-0.546,-0.016]* | -0.14 | [-0.308,0.029] | -0.137 | [-0.282,0.000] |
| incong/spatial | $\mu_{run1}$ | 1.35 | [0.912,1.875] | 2.01 | [1.664,2.408] | 2.54 | [2.190,2.912] |
| | $\mu_{run2}$ | 0.96 | [0.629,1.347] | 1.23 | [0.943,1.578] | 2.05 | [1.568,2.593] |
| | $\mu_{run3}$ | 0.7 | [0.456,0.978] | 0.9 | [0.674,1.160] | 1.53 | [1.166,1.924] |
| | $\mu_{run4}$ | 0.8 | [0.534,1.100] | 0.82 | [0.640,1.018] | 1.58 | [1.266,1.915] |
| | $\mu_{run5}$ | 0.74 | [0.516,1.002] | 0.77 | [0.594,0.966] | 1.55 | [1.217,1.918] |
| | $\mu_{run6}$ | 0.77 | [0.489,1.113] | 0.79 | [0.611,0.994] | 1.37 | [1.032,1.782] |

| | | | | | | |
|---|---|---|---|---|---|---|
| $\mu_{all\ runs}$ | 0.874 | [0.749,1.007] | 1.05 | [0.953,1.150] | 1.74 | [1.586,1.901] |
| $\beta_{lin}$ | -0.084 | [-0.157,-0.008]* | -0.163 | [-0.210,-0.115]* | -0.123 | [-0.178,-0.066]* |
| $\beta_{log}$ | -0.264 | [-0.567,0.022] | -0.377 | [-0.573,-0.185]* | -0.206 | [-0.421,0.007] |

*Notes: non=nonplanner; heur=heuristic; cong=congruent cursor; incong=incongruent cursor; spatial=cursor-normalised spatial skill; coll.=collapsed across runs; runwise and collapsed HDIs have been re-adjusted (subtraction of -1) to discount the constant added to all trials prior to fitting; *=time-on-task coefficient credibly departs 0.*

*Cohort-specific DDM group classifications*

To test group allocations from the DDM for each cohort, we fitted a summary Bayesian multinomial model. The model used a k=3 multinomial likelihood function to characterise the counts (#) for each group classification y=[#(route) #(heuristic) #(nonplanner)], separately for the participants ($n_s$) each cohort (s) $y_s$~Multinomial($\Theta_s$,$n_s$). We assigned $\Theta_s$ an uninformed prior from a Dirichlet distribution $\Theta_s$~Dirichlet($\alpha$=[1,1,1]). Results from this model (summarised in Supplementary Table 3 below) confirmed the DDM ascribed similar group allocations for both cohorts.

**Supplementary Table 4:** cohort-specific DDM group allocations

| | cohort 1 | | cohort 2 | |
|---|---|---|---|---|
| | $\mathbb{E}$(x) | HDI(x) | $\mathbb{E}$(x) | HDI(x) |
| Θ p(route) | 0.370 | [0.167,0.569] | 0.350 | [0.213,0.493] |
| Θ p(heuristic) | 0.210 | [0.054,0.386] | 0.300 | [0.164,0.435] |
| Θ p(nonplanner) | 0.421 | [0.220,0.633] | 0.350 | [0.215,0.492] |

*Acceleration dynamics*

At a resolution of 60 Hz, cursor position during action execution is updated for each frame f by adding a two-element vector ($\mathbb{D}$) to the cursor's position at frame f-1. $\mathbb{D}$ is computed using $D = v = 1$⍰$3$⍰ $\mathbb{P}$⍰$v$⍰ $* \mathbb{V}(v)$⍰. Here, $\mathbb{P}_v$ is the two-element vector (x,y) in screen coordinates describing a Euclidean displacement of 0.320 ° in the direction of a given throttle (v). $\mathbb{V}$(v) scales each coordinate in $\mathbb{P}_v$ in accordance with nonlinear acceleration by using $\mathbb{V}(\mathrm{v}) = f(\mathbb{T}(\mathrm{v}))$. Where$f(x) = -0.011\ x$⍰$3$⍰ $+ 0.167\ x$⍰$2$⍰. For every frame a given throttle (v) is down, the relevant element of three-element vector $\mathbb{T}$ (i.e., $\mathbb{T}$(v)) increases by 0.017 s, and for every frame a throttle is released, $\mathbb{T}$(v) decreases by 0.017 s until it reaches 0. Elements of $\mathbb{T}$ therefore update separately and gradually at this fixed rate, meaning nonzero momentum from one vector can continue influencing the displacement of the cursor after its release and while another throttle is down, allowing curvilinear two-dimensional displacement (see top panel of Figure 1f). However, if more than one throttle is down for a given frame, each element of $\mathbb{T}$ decreases by 0.017 s (unless already at 0), precluding participants from using simultaneous throttle pulsing to create additional displacement angles outside of the six afforded across the two cursors.

*Optimal route simulations*

To enumerate action values derived from route planning we first computed forward simulations of the optimal routes (i.e., with the highest reward yield) from S to G for each cursor on each trial. Separately for each cursor, we first assessed whether the SG on each trial afforded a single linear displacement with one of its vectors from S that would intersect the circular threshold around G (point of intersection=$G^*$). If a cursor satisfied this requirement we computed the optimal throttle sequence with that vector as a single pulse of length $t_{opt}$ that accelerated the cursor to a maximum speed at half the distance between S and $G^*$, followed by a release of the throttle to allow the cursor's momentum to bring it to $G^*$, arriving at a velocity of 0. $t_{opt}$ is estimated to the precision of our (60 Hz) screen resolution by finding the lowest number of frames (λ), such that: $0□\lambda□$ $D□\lambda□ * □ > D/2□$, where D is the Euclidean distance between S and $G^*$ in screen coordinates and $D□\lambda□ * □ = □ SS(\mathbb{P}□v□ * f(\lambda * 0.017))□$, where SS denotes sum of squares and $f(x)$ and $\mathbb{P}_v$ are from the above section describing task physics. Expressing optimal pulse length ($t_{opt}$) in frames (λ) automatically computes the number of units of fuel depleted by this optimal sequence. We subtract λ from 360 as our final estimate of the reward obtainable from the optimal route. (Note that we leave this score on a scale of 0 to 360 for modeling purposes, but present score feedback to participants on each trial as a more intuitive proportion of preserved fuel).

If a cursor does not provide a single linear displacement solution, its optimal route instead comprises a two-pulse sequence using its two vectors that most closely align with the trajectory of the SG, i.e., the two vectors ($v_1$ and $v_2$) with the smallest "offset" values ($\theta_1$ and $\theta_2$) as computed in Figure 2b. The shortest combined displacement of these two vectors that moves a cursor from S to its most nearby Euclidean point on the circular threshold around G ($G^{**}$) can be computed by first originating $v_1$ at S and $v_2$ at $G^{**}$ and finding where they intersect (∩). Forming an oblique triangle with lines |S∩|, |∩$G^{**}$| and |$G^{**}$S|, the length of |S∩| and |∩$G^{**}$| (i.e., the singular displacements of $v_1$ and $v_2$) can then be solved using the law of sines, i.e., $|S \cap| = |G□ ** □S| * sin(\theta□2□)/sin(60)$ and $|\cap G□ ** □| = |G□ ** □S| * sin(\theta□1□)/sin(60)$. Optimal throttle sequence with these vectors is a vector of pulses ($T_{opt}$) containing [$t_{v1}$, $t_{v2}$], respectively solved with the lowest [λ1,λ2] values such that $0□\lambda1□\ D□\lambda□ * □ > D□v1□/2□$ and $0□\lambda2□\ D□\lambda□ * □ > D□v2□/2□$, where $D_{v1}$ is the Euclidean distance between S and ∩, and $D_{v2}$ is the Euclidean distance between $\cap$ and $G^{**}$. Given that $\lambda2$ is calculated from 0 velocity , the optimal sequence pulses v2 immediately upon the release of v1. We subtract $\lambda_{total}$ from 360 as our final estimate of the reward obtainable from the optimal route, where $\lambda_{total}=\lambda1+\lambda2$.

In most cases $\lambda_{total}$ is the same value whether using the above order, or by originating $v_2$ at S and $v_1$ at $G^{**}$, and estimating $[t_{v2}, t_{v1}]$ relative to the resulting intersection ($\cap'$). The exception occurs when one intersection ($\cap$ or $\cap'$) falls outside the grid, requiring more than one direction change to avoid catastrophic error with this sequence. However, all trials had at least one sequence with an intersection inside the grid for each cursor, i.e., at least one optimal path involving a single direction change. Our modeling framework simply required the lowest $\lambda_{total}$ for each cursor on each trial, i.e., either $\lambda$ from a single linear displacement, $\lambda_{total}$ for either route if both intersections fall within the grid, or $\lambda_{total}$ corresponding to the route with its intersection inside the grid, if one fell outside it.
